# A new species of *Nanhsiungchelys* (Testudines: Cryptodira: Nanhsiungchelyidae) from the Upper Cretaceous of Nanxiong Basin, China, and the role of anterolateral processes on the carapace in drag reduction

**DOI:** 10.1101/2022.09.16.506868

**Authors:** Yuzheng Ke, Imran A. Rahman, Hanchen Song, Jinfeng Hu, Kecheng Niu, Fenglu Han

## Abstract

Nanhsiungchelyidae are a group of large turtles that lived in Asia and North America during the Cretaceous. Here we report a new species of nanhsiungchelyid, *Nanhsiungchelys yangi* sp. nov., from the Upper Cretaceous of Nanxiong Basin, China. This is the second valid species of *Nanhsiungchelys*, and the holotype consists of a well-preserved skull and lower jaw, as well as the anterior parts of the carapace and plastron. The diagnostic features of *Nanhsiungchelys* include a huge estimated body size (~55.5 cm), a special network of sculptures on the surface of the skull and shell, weak cheek emargination and temporal emargination, deep nuchal emargination, and a pair of anterolateral processes on the carapace. However, *Nanhsiungchelys yangi* differs from the other species of *Nanhsiungchelys* in having a triangular-shaped snout and wide anterolateral processes. A phylogenetic analysis of nanhsiungchelyids places *Nanhsiungchelys yangi* and *Nanhsiungchelys wuchingensis* as sister taxa. Some nanhsiungchelyids bear special anterolateral processes on the carapace, which are unknown in extant turtles. Here we test the function of these processes in *Nanhsiungchelys yangi* using computational fluid dynamics, and the results suggest these processes could enhance locomotory performance by remarkably reducing drag force when the animal was swimming through water.

## Introduction

Nanhsiungchelyidae are an extinct group of Pan-Trionychia, which lived in Asia and North America from the Early Cretaceous until their extinction at the Cretaceous-Paleogene boundary (Hirayama et al., 2000; Li & Tong, 2017; Joyce et al., 2021). These turtles are characterized by a large body size (maximum 111 cm) (Tong & Li, 2019), flat carapace (Brinkman et al., 2015), stubby limbs (Yeh, 1966), and shells covered with a special network of sculptures consisting of pits and ridges (Li & Tong, 2017). In China, five species of nanhsiungchelyids have been reported (Table 1), with many specimens recovered from the Upper Cretaceous of Nanxiong Basin, Guangdong Province. Yeh (1966) described the first species, *Nanhsiungchelys wuchingensis*, which was restudied by Tong & Li (2019). Hirayama et al. (2009) provided a preliminary study of a large Cretaceous turtle (SNHM 1558) which they placed within Nanhsiungchelyidae; Li & Tong (2017) later attributed this to *Nanhsiungchelys*. In addition, two eggs (IVPP V 2789) from Nanxiong Basin were assigned to nanhsiungchelyids based on the co-occurrence with *Nanhsiungchelys wuchingensis* (Young, 1965).

**Table 1.** Taxonomy and distribution of Nanhsiungchelyidae in China

| Taxa | Specimen Number | Location | Age | Stratigraphic Unit | References |
| --- | --- | --- | --- | --- | --- |
| *Nanhsiungchelys wuchingensis* | IVPP V3106 | Nanxiong,<br>Guangdong | Late<br>Cretaceous | Yuanpu<br>Formation | Yeh (1966)<br>Tong & Li (2019) |
| *Nanhsiungchelys* sp. | SNHM 1558 | Nanxiong,<br>Guangdong | Late<br>Cretaceous | Nanxiong Group | Hirayama et al. (2009)<br>Li & Tong (2017) |
| *Nanhsiungchelys yangi* sp. nov. | CUGW VH108 | Nanxiong,<br>Guangdong | Late<br>Cretaceous | Yuanpu<br>Formation | This paper |
| *Jiangxichelys neimongolensis* | IVPP RV96007, IVPP<br>RV96008, IVPP<br>290690-6 RV 96009,<br>IVPP 020790-4 RV<br>96010, IVPP 130790<br>RV 96011, IMM 4252,<br>IMM 2802, IMM<br>96NMBY-I-14, IMM<br>93NMBY-2 | Bayan<br>Mandahu,<br>Inner<br>Mongolia | Late<br>Cretaceous | Wulansuhai<br>Formation | Brinkman & Peng (1996)<br>Brinkman et al. (2015)<br>Li & Tong (2017) |
| *Jiangxichelys ganzhouensis* | NHMG 010415,<br>JXGZ(2012)-178,<br>JXGZ(2012)-179,<br>JXGZ(2012)-180,<br>JXGZ(2012)-182 | Ganzhou,<br>Jiangxi | Late<br>Cretaceous | Nanxiong<br>Formation | Tong & Mo (2010)<br>Tong et al. (2016) |
| *Yuchelys nanyangensis* | HGM NR09-11-14,<br>CUGW EH051 | Nanyang,<br>Henan | Late<br>Cretaceous | Xiaguan<br>Formation | Tong et al. (2012)<br>Ke et al. (2021) |

Recently, the phylogenetic relationships of nanhsiungchelyids have been studied in detail (Danilov et al., 2013; Brinkman et al., 2015; Tong et al., 2016; Mallon & Brinkman, 2018; Tong & Li, 2019). Among the 13 species of Nanhsiungchelyidae, *Nanhsiungchelys* and *Anomalochelys* form a sister group, which share an elongated shell, huge nuchal emargination, large anterior process on the carapace, wide neurals and vertebral scutes, and sub-triangular first vertebral with very narrow anterior end (Tong & Li, 2019). The other nanhsiungchelyids (*Basilemys*, *Hanbogdemys*, *Jiangxichelys*, *Kharakhutulia*, *Yuchelys*, and *Zangerlia*) usually have a shorter carapace, shallow nuchal emargination, narrow neurals and vertebral scutes, and lack large anterior processes on the carapace (Tong & Li, 2019). *Nanhsiungchelys* and *Anomalochelys* have only been found in southern China and Japan (Hirayama et al., 2001; Hirayama et al., 2009; Li & Tong, 2017; Tong & Li, 2019), but the other species have a wider distribution (Central Asia, East Asia, and North America) (Danilov & Syromyatnikova, 2008; Mallon & Brinkman, 2018).

The ecology of nanhsiungchelyids is debated (Mallon & Brinkman (2018) provided a detailed overview). Although many researchers support a terrestrial mode of life for the group (Yeh, 1966; Hutchison & Archibald, 1986; Scheyer, 2007; Dudgeon et al., 2021), Sukhanov & Narmandakh (1977) argued that the anatomy of *Hanbogdemys* was inconsistent with terrestrial habits based on characteristics of the forelimbs, humerus, and pelvis, and Nessov (1981) regarded these animals as specialized swimmers. Moreover, nanhsiungchelyids usually have a flatted carapace, which is a common feature of aquatic turtles (Xiao et al., 2017). Some nanhsiungchelyids (i.e. *Nanhsiungchelys* and *Anomalochelys*) have distinctive anterolateral processes on the carapace (Hirayama et al., 2001; Hirayama et al., 2009; Tong & Li, 2019). The main function of these processes was previously thought to be protection (Hirayama et al., 2001), but their ecological significance has not received much attention from researchers to date. Although these anterolateral processes have not been found in any extant turtles, similar horn-like structures at the anterolateral margin of the carapace occurred in the Miocene turtle *Stupendemys geographicus*, which is thought to be an aquatic side-necked turtle (Cadena et al., 2020). Further support for an aquatic existence comes from *Anomalochelys angulata*, which was recovered from marine sediments containing numerous radiolarian fossils (Hirayama et al., 2001), indicating that they lived in a coastal environment. Together, this strongly suggests that nanhsiungchelyids (especially *Nanhsiungchelys* and *Anomalochelys*) were capable of swimming, but further analyses are necessary to test the function of the anterolateral processes in water.

Here, we report a new species of *Nanhsiungchelys* from Nanxiong Basin based on a complete skull and partial postcranial skeleton. This allows us to explore the taxonomy and morphology of nanhsiungchelyids, and based on this we carry out a phylogenetic analysis of the group. In addition, we use computer simulations of fluid flow (computational fluid dynamics) to obtain new insights into the function of the anterolateral processes.

### Geological Setting

Nanxiong Basin is a NE-trending faulted basin controlled by the Nanxiong Fault in the northern margin, covering an area of about 1800 km^2^ and spanning Guangdong and Jiangxi provinces in China (Zhang et al., 2013). There are well-exposed outcrops of Cretaceous–Paleogene strata in Nanxiong Basin (Ling et al., 2005), which consist of nine formations: the Cretaceous Changba Formation, Jiangtou Formation, Yuanpu Formation, Dafeng Formation, Zhutian Formation, Zhenshui Formation, and Shanghu Formation (Pingling Member); and the Palaeocene Shanghu Formation (Xiahui Member), Nongshan Formation, and Guchengcun Formation (Zhang et al., 2013). Of these, the first six formations (Changba Formation to Zhenshui Formation) are referred to as the Nanxiong Group (Chang & Tung, 1963; Zhang et al., 2013). The Yuanpu Formation comprises a set of fine-grained sedimentary strata, which are interbedded with brownish red to purplish red thick-bedded siltstones and light brown to maroon medium thick-bedded sandstones of unequal thickness, and is locally intercalated with sandy conglomerate and a thin layer of gravel-bearing sandstone or lens (Zhang et al., 2013). Zhao et al. (1991) reported two K–Ar ages for the Yuanpu Formation (67.04±2.31 Ma and 67.37±1.49 Ma), which indicate that it was deposited in the late Maastrichtian stage. Many vertebrate fossils have been recovered from Yuanpu Formation, including: the turtle *Nanhsiungchelys wuchingensis* (Yeh, 1966; Tong & Li, 2019); the turtle eggs *Oolithes nanhsiungensis* (Young, 1965); and the dinosaur eggs *Macroolithus rugustus*, *Nanhsiungoolithus chuetienensis*, *Ovaloolithus shitangensis*, *Ovaloolithus nanxiongensis*, and *Shixingoolithus erbeni* (Zhao et al., 2015).

## Material and Method

### Fossil specimen

The specimen (CUGW VH108) consists of a well-preserved skull and lower jaw, together with the anterior parts of the carapace and plastron (Figures 1–3). It is housed in the paleontological collections of China University of Geosciences (Wuhan). The skeleton was prepared by Yuzheng Ke and Kaifeng Wu, using an Engraving Pen AT-310, and was photographed with a Canon EOS 6D camera.

**Figure 1.**
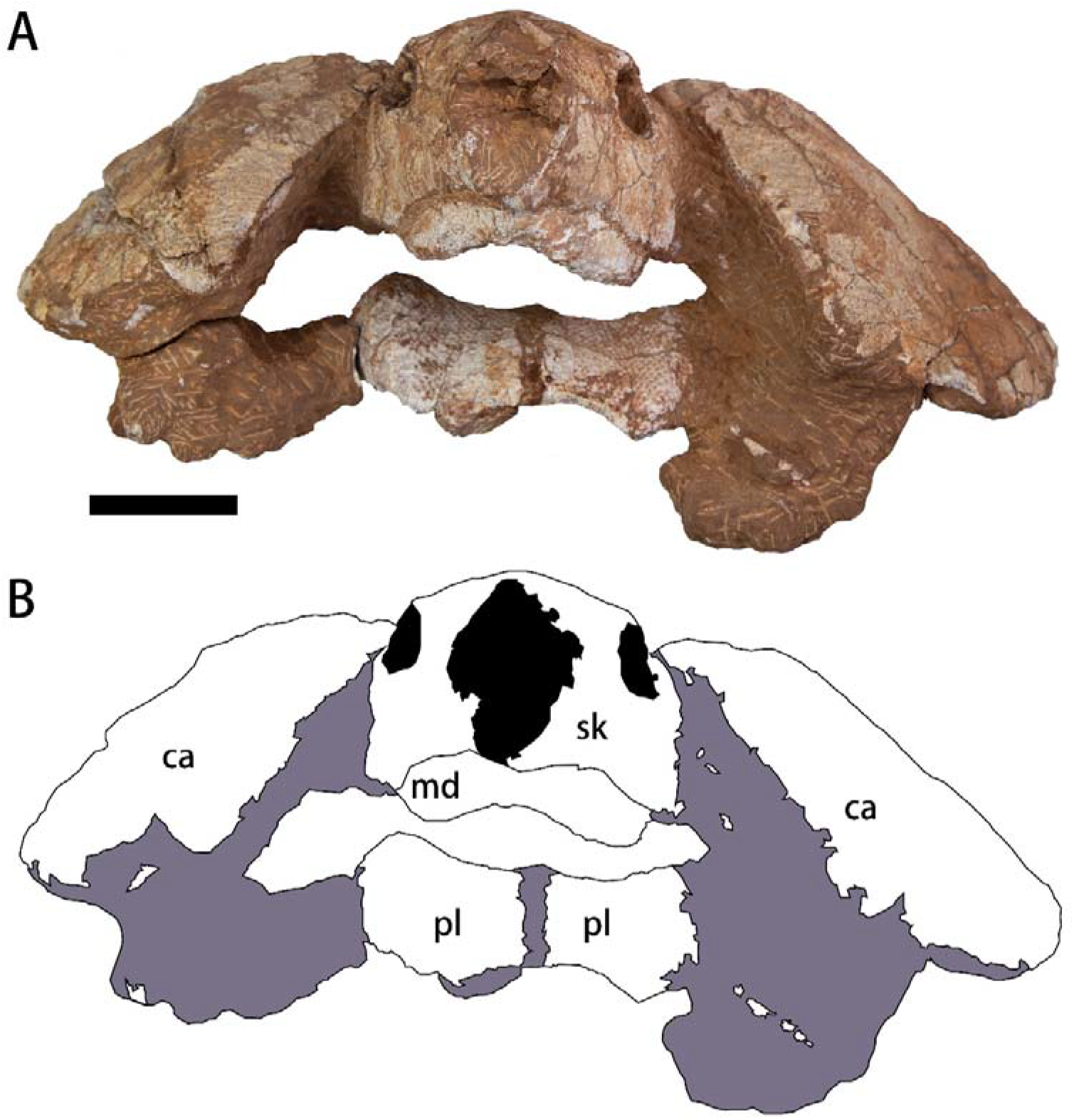
Photograph (A) and outline drawing (B) of *Nanhsiungchelys yangi* (CUGW VH108) in anterior view. Gray and black parts indicate the surrounding rock and openings of the skull, respectively. Scale bar equals 5 cm. Abbreviations: ca, carapace; md, mandible; pl, plastron; sk, skull.

**Figure 2.**
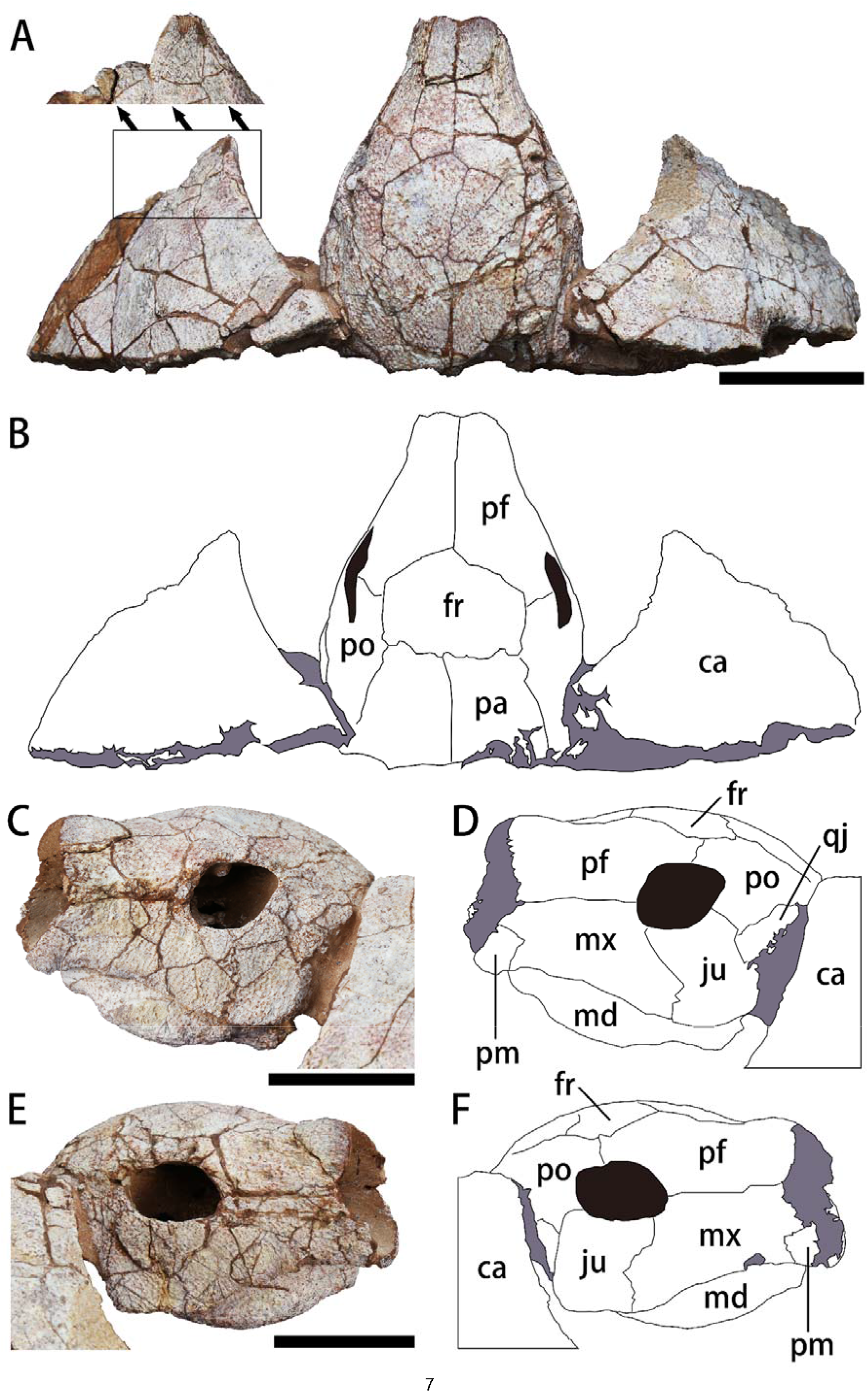
The skull and carapace of *Nanhsiungchelys yangi* (CUGW VH108). A, B. Photograph and outline drawing of the skull and carapace in dorsal view, and the small figure shows details in black box (perpendicular to surface of carapace). C, D. Photograph and outline drawing of the skull in left lateral view. E, F. Photograph and outline drawing of the skull in right lateral view. Gray and black parts indicate the surrounding rock and openings of the skull, respectively. Scale bars equal 5 cm. Abbreviations: ca, carapace; fr, frontal; ju, jugal; md, mandible; mx, maxilla; pf, prefrontal; pa, parietal; pm, premaxilla; po, postorbital; qj, quadratojugal.

**Figure 3.**
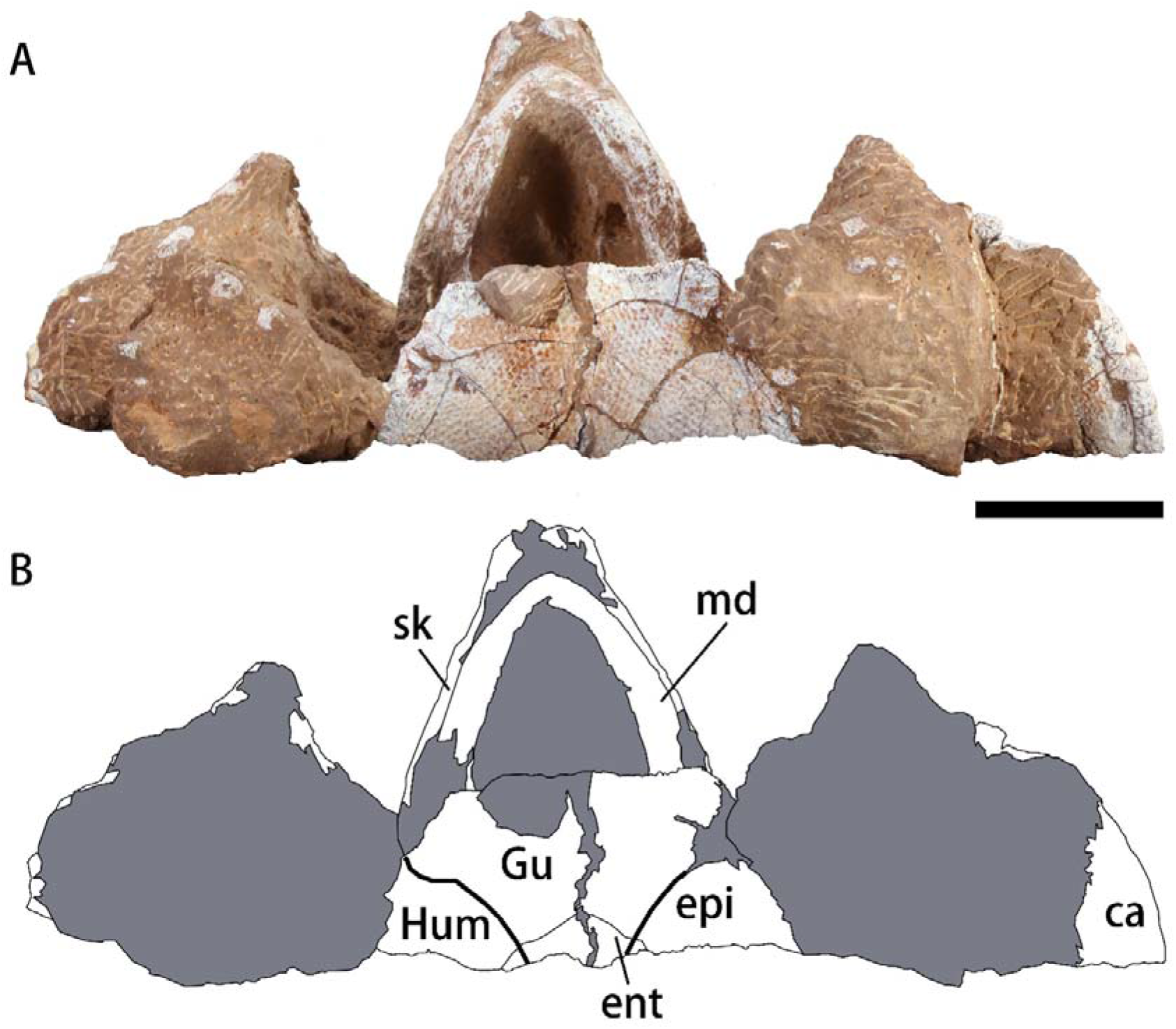
Photograph (A) and outline drawing (B) of *Nanhsiungchelys yangi* (CUGW VH108) in ventral view. Bold lines represent the sulci between scutes and gray parts indicate the surrounding rock. Scale bar equals 5 cm. Abbreviations: ca, carapace; epi, epiplastron; ent, entoplastron; Gu, gular scute; Hum, humeral scute; md, mandible; sk, skull.

### Phylogenetic analysis

Parsimony phylogenetic analysis was performed using the software TNT 1.5 (Goloboff & Catalano, 2016). The data matrix used herein was updated from Tong & Li (2019) and Mallon & Brinkman (2018), and includes 17 taxa and 50 characters. Because there are five inframarginal scutes on *Jiangxichelys ganzhouensis* (Tong et al., 2016), character 37 was modified to: “Inframarginals: (0) five to three pairs; (1) two pairs; (2) absent”. In addition, character 48 was changed in *Jiangxichelys ganzhouensis* from ? to 1 (i.e. ratio of midline epiplastral suture length to total midline plastral length greater than 0.1). A new character was also added: ratio of length to width of the carapace: (0) less than 1.6; (1) equal to or larger than 1.6. Moreover, *Yuchelys nanyangensis* was added to the data matrix based on Tong et al. (2012). A total of 13 characters out of 50 could be coded for *Nanhsiungchelys yangi*, representing only 26 % of the total number of characters. This is because the new species is based on a partial specimen missing many of the features scored in other taxa. The analysis was conducted using a traditional search with 1000 replicates. A tree bisection reconnection (TBR) swapping algorithm was employed, and 10 trees were saved per replicate. Most characters were treated as unordered, but characters 24, 29, and 47 were set to be additive because they show continuous character changes (i.e. 0→2). All characters are of equal weight. Standard bootstrap support values were calculated using a traditional search with 100 replicats. Bremer support values were also calculated (Bremer, 1994).

### Computational fluid dynamics

Computational fluid dynamics (CFD) is an useful tool for simulating flows of fluids and their interaction with solid surface (Rahman, 2017; Gibson et al., 2021). This basic principle involves transforming the Navier–Stokes equations corresponding to flow problems into algebraic equations and solving them using certain numerical methods at finite discrete moments and spatial nodes (grids) (Guo et al., 2019). Recently, CFD techniques have been used in paleontology to quantitatively assess the habits and ecology of a wide range of extinct organisms (Shiino et al., 2009; Shiino & Kuwazuru, 2010; Shiino et al., 2012; Kogan et al., 2015; Liu et al., 2015; Dynowski et al., 2016; Gutarra et al., 2019; Rahman et al., 2020; Gibson et al., 2021; Song et al., 2021). In our research, CFD was used to evaluate the drag forces of turtles in water, and the simulations were performed in the software COMSOL Multiphysics (v. 5.6).

#### Digital modelling

Considering the close relationship between *Nanhsiungchelys yangi* and *Nanhsiungchelys wuchingensis* in our phylogeny (see below), we reconstructed a full 3-D model of *Nanhsiungchelys yangi* (Figure 4A–C) by referring to the holotype of *Nanhsiungchelys wuchingensis* (IVPP V3106). The 3-D reconstruction was created using the in-built geometry tools in COMSOL. The main structures of the turtle model were created with simple shapes (e.g. ellipsoids and cylinders). In addition, interpolation curves were drawn in Plane Geometry, which were then further extended into faces. Lastly, several fillets were added to create rounded corners on the 3-D geometries. This model was scaled to a carapace length of 1.0 m based on well-preserved specimens of *Nanhsiungchelys wuchingensis* (IVPP V3106) and *Nanhsiungchelys* sp. (SNHM 1558) which are 0.87 to 1.11 m in total carapace length (Hirayama et al., 2009; Tong & Li, 2019). Considering extant turtles swim with their heads and necks stretching from the shells, an idealized head and neck were added to the model to give the modal a total length of 1.25 m. In addition, we constructed an idealized 3-D model of a turtle without the anterolateral processes on the carapace (here referred to as ‘generalized turtle’) for comparison, whose total length was also 1.25 m (Figure 4D–F).

**Figure 4.**
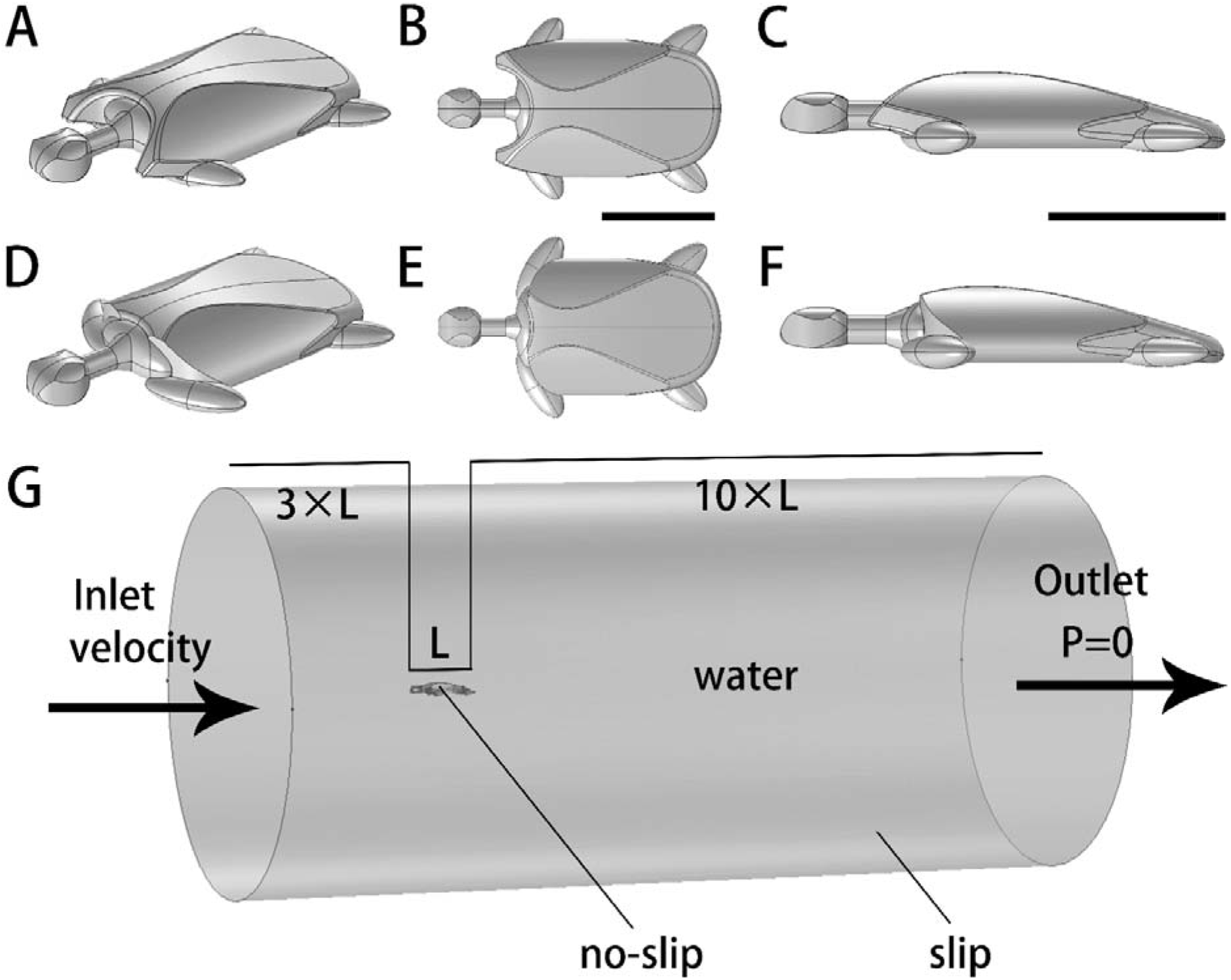
Three-dimensional digital models of *Nanhsiungchelys yangi* (A–C), a generalized turtle (D–F), and the computational domain used for computational fluid dynamics simulations (G). Scale bars equal 0.5 m.

For each model of length *L*, a cylindrical computational domain was created, whose upstream length was 3×*L*, downstream length was 10×*L*, and radius was 5× the maximum width of the model, following Gutarra et al. (2019) (Figure 4G). As these models are bilaterally symmetrical, only half of the turtle models and cylinders were used in simulations in order to reduce the computation time.

#### Fluid properties and boundary conditions

Parts of the domain inside the cylinder, surrounding the turtle model, were assigned the material properties of water using the built-in materials library in COMSOL. The upstream end of the cylinder was set as the inlet (turbulent intensity is 0.05, with the flow velocity specified here) and the downstream end of the cylinder was set as the outlet (pressure condition is static, pressure specified as 0 Pa, and suppress backflow is selected) (Figure 4G). The swimming speeds of extinct nanhsiungchelyids are unknown; however, the modal and maximum swimming speeds of the extant leatherback sea turtle *Dermochelys coriacea* (which has a curved carapace length from 1.45 to 1.69 m) are known to range from 0.56–0.84 m/s and 1.9–2.8 m/s, respectively (Eckert, 2002). It is highly unlikely the swimming speeds of nanhsiungchelyids were faster than leatherback sea turtles due to the lack of paddle-like limbs, and we therefore simulated swimming speeds of 0.6 m/s, 1.0 m/s, 1.4 m/s, 1.8 m/s, 2.2 m/s, and 2.6 m/s in our study.

The flow regime was characterized using the dimensionless Reynolds numbers (*Re*) (Reynolds, 1883; Gibson et al., 2021):

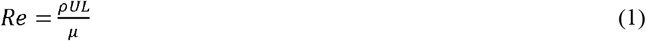

where *ρ* is the density of water (1000 kg/m^3^), *U* is the velocity of water flow (m/s), *L* is the model’s maximum width (m), and the *μ* is water’s dynamic-viscosity coefficient. The *Re* of our simulations was ~4.75×10^5^ to ~2.06×10^6^, which falls within the range of turbulent flow (i.e. *Re* > 1×10^4^) (Gutarra et al. (2019), supplementary information). As a result, the *k*-*ε* turbulence model was used in all our simulations, which is robust, economizes on computational cost, and is known to be reasonably accurate for a wide range of turbulent flows (Adkins & Yan, 2006). Slip boundary condition was assigned to the side of the cylindrical computational domain, and no-slip boundary condition was assigned to the surface of the 3-D turtle models.

#### Mesh size and computation

The domains were meshed using free tetrahedral elements, with prismatic boundary layer elements inserted along the interface between the turtle model (Figure 5). A stationary solver was used to compute the steady state flow patterns, with the segregated iterations terminated when the relative tolerance reached 1×10^−4^.

**Figure 5.**
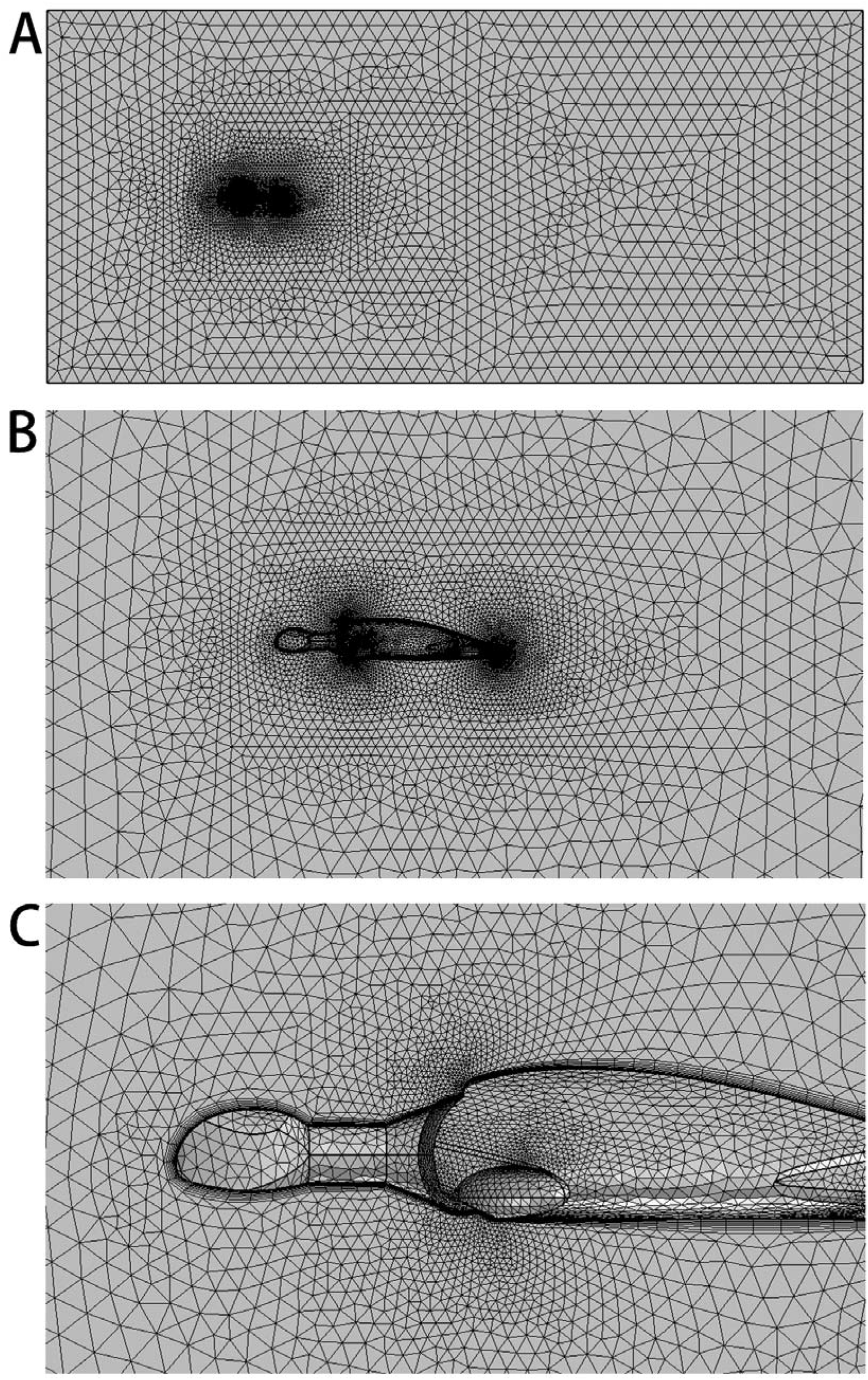
Mesh used in CFD simulations. A, mesh used in the whole computational domain. B, details of the mesh near the turtle model. C, details of the mesh near the anterior of the turtle model.

To evaluate the effect of mesh size on the CFD results, sensitivity tests were conducted with different meshes. Using an inlet velocity of 1.0 m/s, three different mesh sizes (‘normal’ to ‘finer’) were used for each of the models, and this showed that the drag forces and drag coefficients did not change significantly (Table 2, Figure 6). As a result, the finer mesh was selected and used in our analyses, which was composed of numerous smaller cells and thus could most accurately represent the flow (Gibson et al., 2021; Rahman, 2017).

**Table 2.** Drag forces and drag coefficients obtained for different mesh sizes for three-dimensional digital models of *Nanhsiungchelys yangi* and a generalized turtle.

|  | *Nanhsiungchelys yangi* |  |  | Generalized turtle |  |  |
| --- | --- | --- | --- | --- | --- | --- |
| Mesh size | Number of elements | Drag force | Drag coefficient | Number of elements | Drag force | Drag coefficient |
| Normal | 380495 | 22.312 N | 0.372 | 316015 | 28.058 N | 0.468 |
| Fine | 644382 | 20.892 N | 0.348 | 495692 | 27.088 N | 0.451 |
| Finer | 1431064 | 19.176 N | 0.320 | 1006082 | 25.826 N | 0.430 |

**Figure 6.**
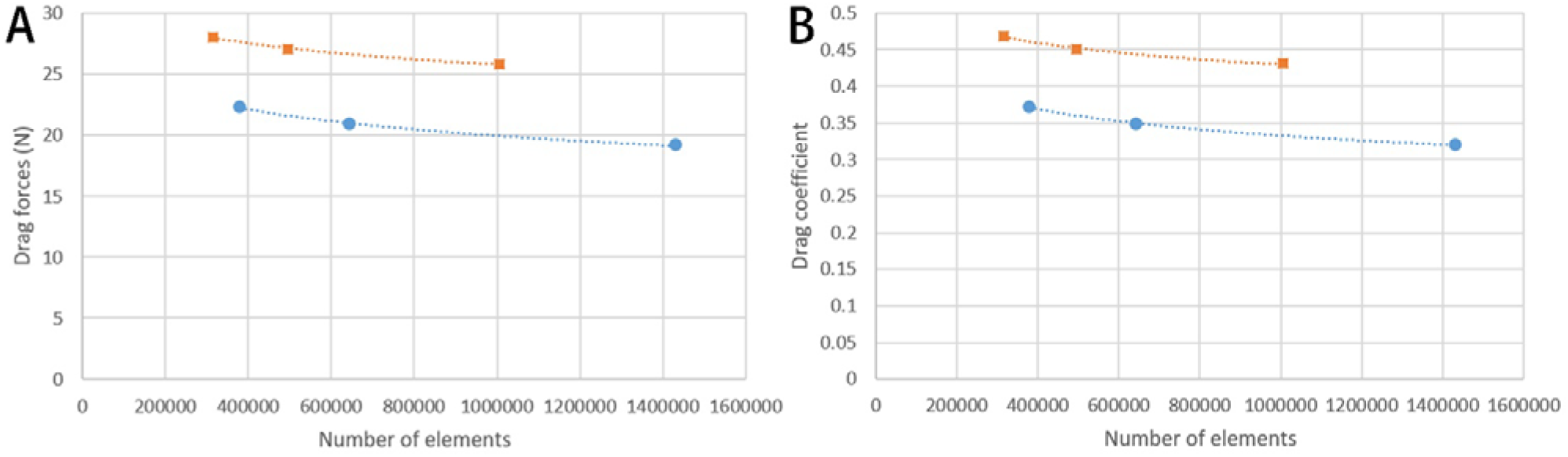
Comparison of drag forces (A) and drag coefficients (B) for three-dimensional digital models of *Nanhsiungchelys yangi* and generalized turtle at different mesh sizes. Blue circles represent results for the *Nanhsiungchelys yangi* model and orange squares represent results for the generalized turtle model.

Drag forces were computed for each model based on surface integration. Drag coefficients (*C_D_*) were then calculated using the following equation (Rahman, 2017; Gibson et al., 2021):

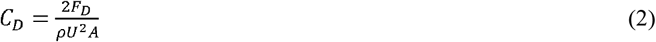

where *F*_*D*_ is the drag force (N), *ρ* is the density of water (1000 kg/m^3^), *U* is the velocity of water flow (m/s), and *A* is the characteristic area (m^2^). Moreover, 2-D plots showing flow velocity magnitude (Figure 10) and streamlines around the turtle models (Figure 11) were visualized.

**Figure 7.**
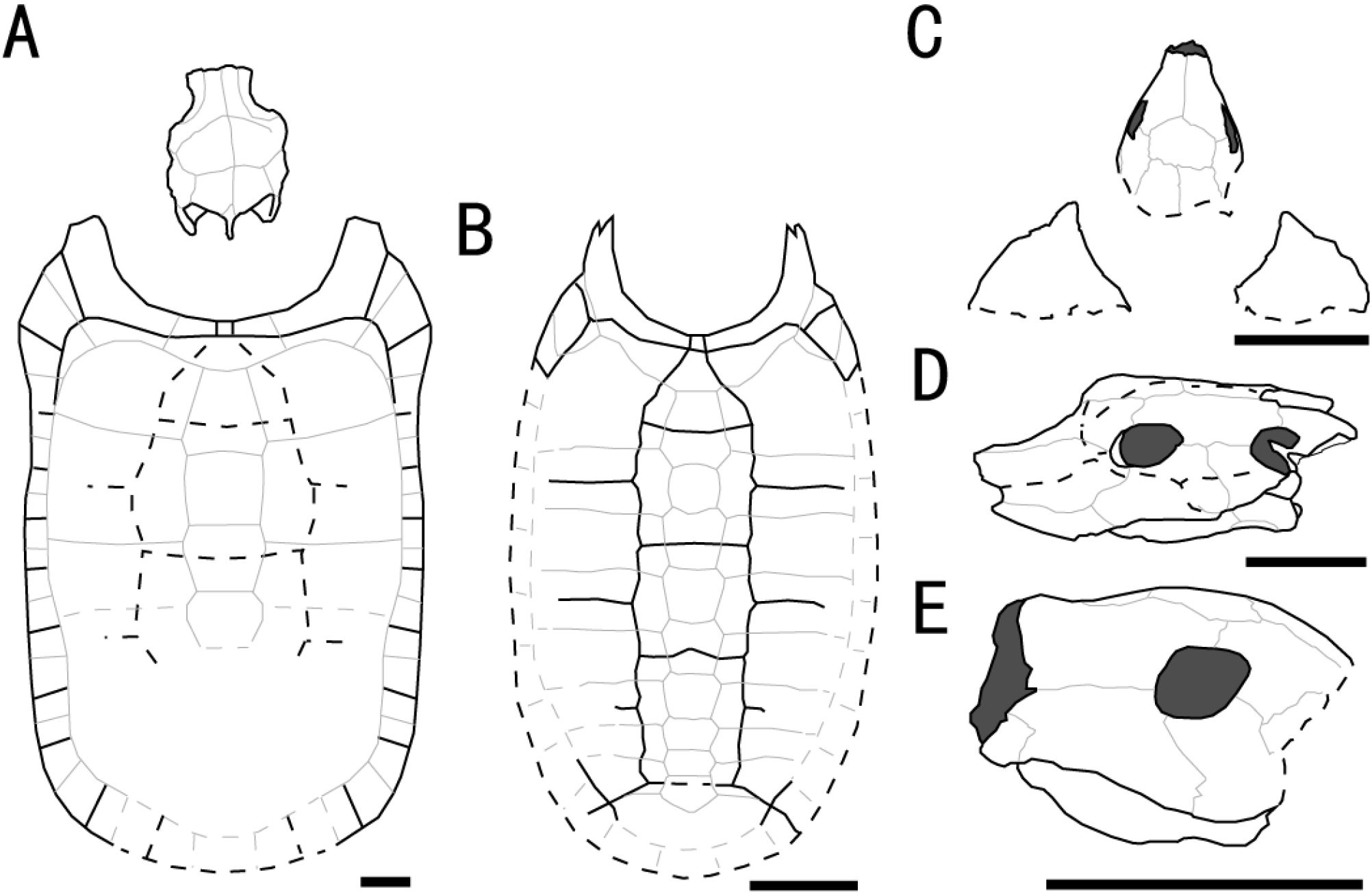
Outline drawings of three nanhsiungchelyids. A. Skull and carapace of *Nanhsiungchelys wuchingensis*, after Hirayama et al. (2001) and Tong & Li (2019). B. Carapace of *Anomalochelys angulata*, after Hirayama et al. (2001). C. Skull and partial carapace of *Nanhsiungchelys yangi* (CUGW VH108). D. Skull of *Nanhsiungchelys wuchingensis* in left lateral view, after Tong & Li (2019). E. Skull of *Nanhsiungchelys yangi* (CUGW VH108) in left lateral view. Scale bars equal 10 cm. Bold black lines represent the sulci between scutes, thin gray lines indicate the sutures between bones, and dashed lines indicate a reconstruction of poorly preserved areas.

**Figure 8.**
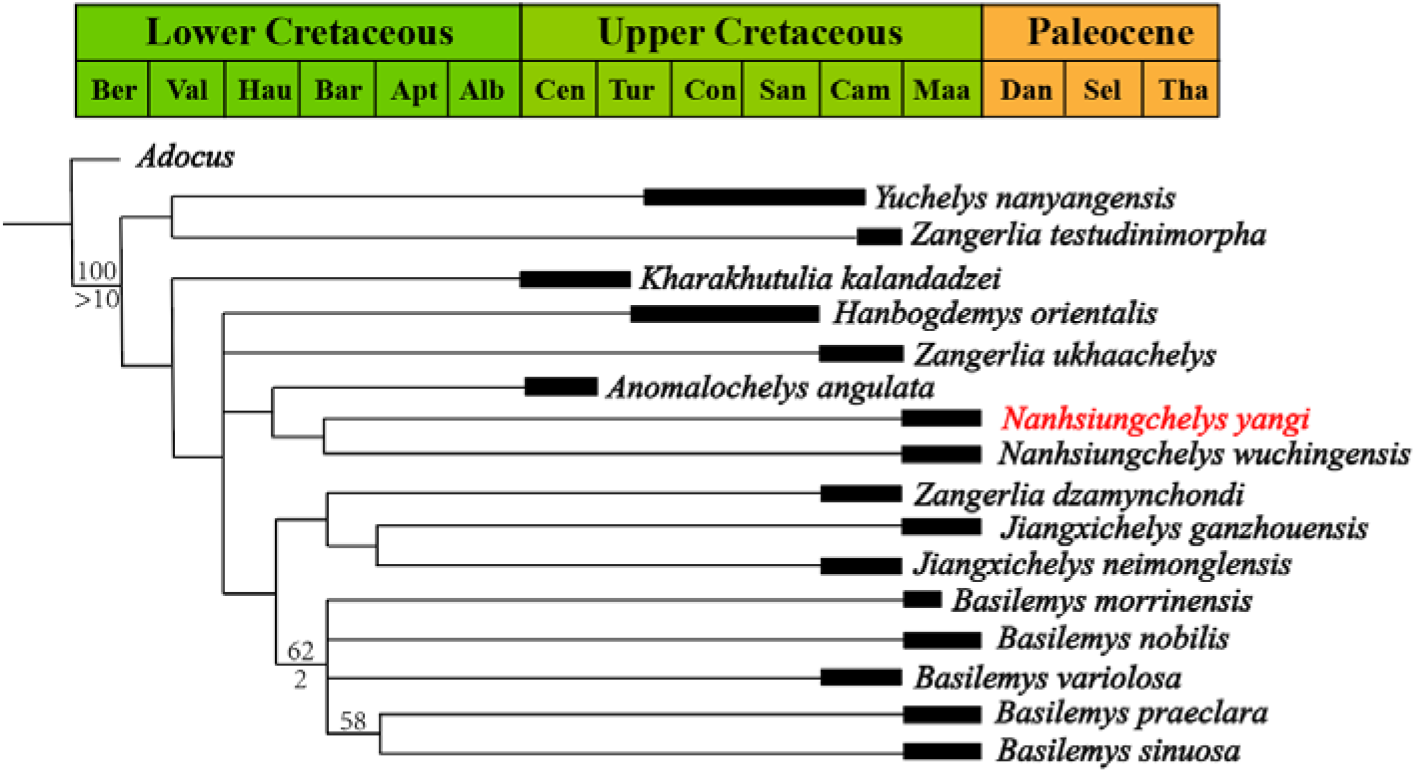
Strict consensus tree of Nanhsiungchelyidae. Numbers above nodes are bootstrap support value, and numbers below nodes are Bremer support values. Temporal distributions of the above species are based on Yu et al. (1990), Danilov et al. (2013), Wang et al. (2013), Zhang et al. (2013), and Mallon & Brinkman (2018). Abbreviations: Ber, Berriasian; Val, Valanginian; Hau, Hauterivian; Bar, Barremian; Apt, Aptian; Alb, Albian; Cen, Cenomanian; Tur, Turonian; Con, Coniacian; San, Santonian; Cam, Campanian; Maa, Maastrichtian; Dan, Danian; Sel, Selandian; Tha, Thanetian.

**Figure 9.**
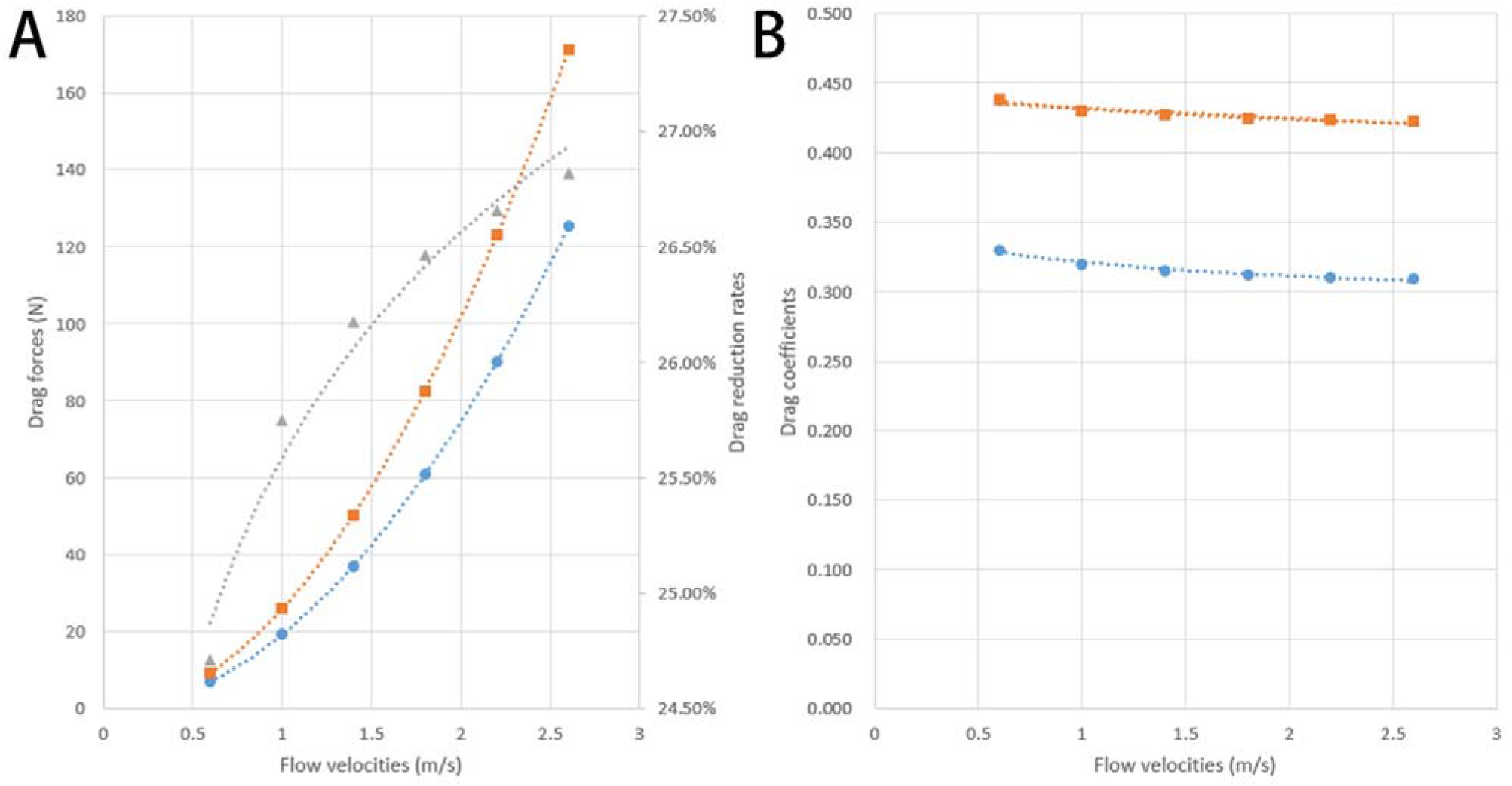
Comparison of drag forces (A) and drag coefficients (B) for three-dimensional digital models of *Nanhsiungchelys yangi* and a generalized turtle at different flow velocities. Blue circles represent results for the *Nanhsiungchelys yangi* model, orange squares represent results for the generalized turtle model, and grey triangles represent the drag reduction brought about by the anterolateral processes.

**Figure 10.**
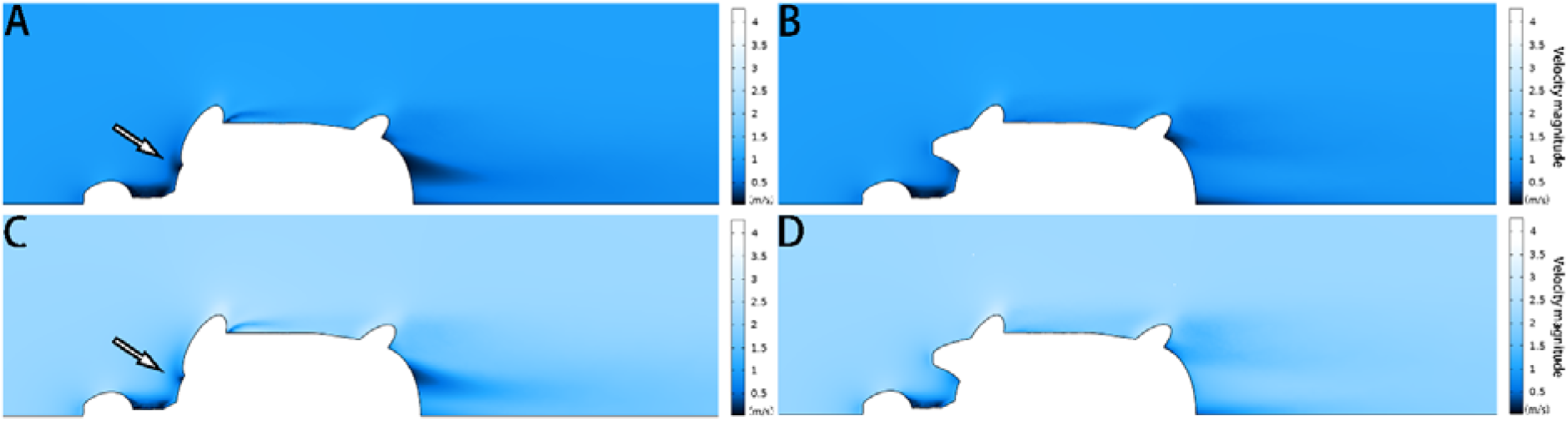
2-D plots of flow velocity magnitude. A shows results for the generalized turtle model at a flow velocity of 1.0 m/s. B shows results for the *Nanhsiungchelys yangi* model at a flow velocity of 1.0 m/s. C shows results for the generalized turtle model at a flow velocity of 2.2 m/s. D shows results for the *Nanhsiungchelys yangi* model at a flow velocity of 2.2 m/s. The arrows indicate the low-velocity zones near the proximal forelimbs.

**Figure 11.**
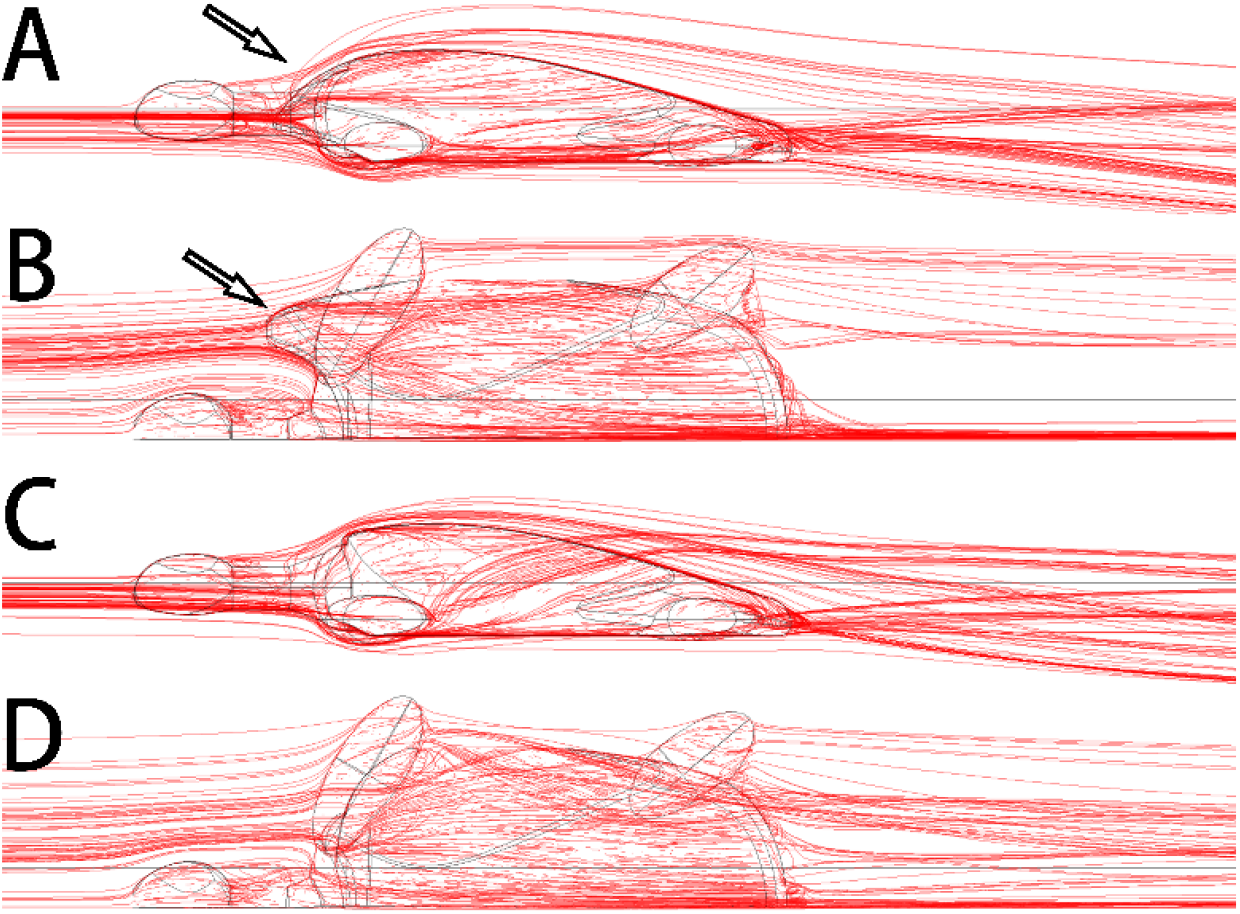
3-D plots of streamline at flow velocities of 1.0 m/s. A. *Nanhsiungchelys yangi* model in left lateral view. B. *Nanhsiungchelys yangi* model in dorsal view. C. Generalized turtle model in left lateral view. D. Generalized turtle model in dorsal view. The arrows indicate the anterolateral processes.

#### Institutional abbreviations

CUGW, China University of Geosciences (Wuhan); HGM, Henan Geological Museum; IMM, Inner Mongolia Museum; IVPP, Institute of Vertebrate Paleontology and Paleoanthropology, Chinese Academy of Sciences; LJU, Lanzhou Jiaotong University; NHMG, Natural History Museum of Guangxi; NMBY, Nei Mongo Bowuguan; SNHM, Shanghai Natural History Museum; UB, University of Bristol; UPC, China University of Petroleum (East China); YSNHM, Yingliang Stone Natural History Museum.

## Results

### Systematic paleontology

Order Testudines Linnaeus, 1758

Infraorder Cryptodira Cope, 1868

Superfamily Trionychoidea Fitzinger, 1826

Family Nanhsiungchelyidae Yeh, 1966

Genus *Nanhsiungchelys* Yeh, 1966

#### Emended diagnosis

A genus of Nanhsiungchelyidae of medium-large size, with a carapace length of 0.5–1.1 m. The surface of the skull, lower jaw and both carapace and plastron are covered with sculptures consisting of large pits formed by a network of ridges. Temporal emargination and cheek emargination are weak; orbits located at about mid-length of the skull and facing laterally; jugal forms the lower margin of the orbit. Carapace elongate, with a deep nuchal emargination and a pair of large anterolateral processes that extend forward and are formed entirely by the first peripheral; wide neural plates and vertebral scutes; gulars fused and extend deeply onto the entoplastron; intergulars absent; complete row of narrow inframarginals. Wide angle between the acromion process and scapula process of about 105°. One large dermal plate located above the manus.

#### Type species

*Nanhsiungchelys wuchingensis* Yeh, 1966

#### Distribution

Guangdong, China

Species *Nanhsiungchelys yangi* sp. nov.

#### Etymology

*Yangi* is in memory of paleontologist Zhongjian Yang (Chung-Chien Young).

#### Holotype

CUGW VH108, a partial skeleton comprising a well-preserved skull and lower jaw and the anterior parts of the carapace and plastron (Figures 1–3).

#### Locality and horizon

Nanxiong, Guangdong, China. Yuanpu Formation, Upper Cretaceous.

#### Diagnosis

A medium-sized species of *Nanhsiungchelys* with an estimated carapace length of more than 0.5 meters. It differs from other species of *Nanhsiungchelys* in the following combination of characters: the snout is triangular in dorsal view; the premaxilla is higher than wider; the posteroventral ramus of the maxilla extends to the ventral region of the orbit; the dorsal margin of the maxilla is relatively straight; the jugal is higher than wider; the prefrontal is convex dorsally behind the naris; no suture present between the paired frontals; the temporal emargination is mainly formed by the parietal; the paired parietals are bigger than the fused frontal in dorsal view; the middle and posterior parts of the mandible are more robust than the most anterior part in ventral view; the anterolateral processes is wide; and the angle between the two anterior edges of the entoplastron is wide (~110°).

## Description

### General aspects of the skull

The skull is large, with a length of 13 cm (Figure 2A, B). It is well preserved but there are many cracks on its outer surface, which limit the identification of bone sutures. The snout (i.e. the parts anterior to the orbit) is large, equal to about 1/3 length of the skull, and longer than in *Jiangxichelys neimongolensis* and *Zangerlia ukhaachelys* (Joyce & Norell, 2005; Brinkman et al., 2015). In dorsal view, the snout is triangular and differs from *Nanhsiungchelys wuchingensis* in which the snout is trumpet-shaped (Tong & Li, 2019). A large naris is located in the front part of the snout, which is roughly lozenge shaped and higher than wider in anterior view (Figure 1). Since the skull is partially withdrawn into the shell, it is difficult to accurately determine the morphological characteristics of cheek emargination (Figure 2C–F). However, we infer that the cheek emargination is not deep, because otherwise this would extend beyond the dorsal margin of the orbit, as in some extant turtles (e.g. *Emydura macquarrii*) (Li & Tong, 2017). Posteriorly, the temporal emargination is weakly developed (Figure 2A, B), which is similar to *Nanhsiungchelys wuchingensis* (Tong & Li, 2019), but differs from *Jiangxichelys neimongolensis*, *Jiangxichelys ganzhouensis* and *Zangerlia ukhaachelys* (Brinkman & Peng, 1996; Joyce & Norell, 2005; Tong et al., 2016). The surface of the skull (and the carapace and plastron) is covered with a special network of sculptures consisting of pits and ridges, which is a synapomorphy of Nanhsiungchelyidae (Li & Tong, 2017).

### Premaxilla

A small bone in the anterior and ventral part of the maxilla is identified as the premaxilla (Figure 2C–F). It is higher than wider, similar to *Jiangxichelys neimongolensis* and *Zangerlia ukhaachelys* (Joyce & Norell, 2005; Brinkman et al., 2015), but differs from *Nanhsiungchelys wuchingensis* in which the premaxilla is wider than higher in lateral view and has an inverse Y-shape in ventral view (Tong & Li, 2019). Due to poor preservation of the most anterior parts of the premaxilla, it is difficult to determine whether the left and right premaxillae contact each other.

### Maxilla

The maxilla is large and trapezoid in outline (Figure 2C–F). The main shaft is located anterior to the orbit, but the posteroventral ramus extends to the ventral region of the orbit, which differs from the situation in *Nanhsiungchelys wuchingensis* in which the maxilla is located entirely anterior to the orbit (Tong & Li, 2019), and also differs from most other turtles (including *Zangerlia ukhaachelys* and *Jiangxichelys neimongolensis*) in which the maxilla consists of the lower rim of the orbit (Joyce & Norell, 2005; Brinkman et al., 2015). In lateral view, the dorsal margin of the maxilla is relatively straight and extends posteriorly to the mid-region of the eye socket, which is similar to some extant turtles (e.g. *Platysternon megacephalum*) (Li & Tong, 2017). However, this differs from *Nanhsiungchelys wuchingensis* in which the top of the maxilla is curved dorsally (Tong & Li, 2019), and also differs from *Zangerlia ukhaachelys* and *Jiangxichelys neimongolensis* in which the top of the maxilla tapers anterdorsally (Joyce & Norell, 2005; Brinkman et al., 2015).

### Jugal

The jugal is shaped like a parallelogram in lateral view (Figure 2C–F). It is higher than wider, unlike *Nanhsiungchelys wuchingensis* in which the jugal is wider than higher (Tong & Li, 2019). The jugal consists of the lower rim of the orbit, which is similar to *Nanhsiungchelys wuchingensis*, but differs from most turtles in which this structure is mainly formed by the maxilla (Tong & Li, 2019). The jugal of *Nanhsiungchelys yangi* also differs from *Jiangxichelys ganzhouensis* in which the jugal is more posteriorly located (Tong et al., 2016). The jugal contacts with the maxilla in front, and this boundary is sloped. The terminal parts of the jugal contacts with the quadratojugal.

### Quadratojugal

The bone that behind the jugal and below the postorbital is identified as the quadratojugal (Figure 2C–F). Its location is similar to *Nanhsiungchelys wuchingensis* (Tong & Li, 2019), but the full shape is uncertain due to the cover with carapace.

### Prefrontal

In dorsal view, each prefrontal is large and elongate anteroposterioly, and narrows anteriorly and enlarges posteriorly (Figure 2A, B). The portion in front of the orbit is entirely composed of the prefrontal (Figure 2A, B), which differs from *Nanhsiungchelys wuchingensis* in which the maxilla extends forward to the prefrontal and occupies some space (Tong & Li, 2019). The paired prefrontals contact each other at the midline and form an approximate arrow shape. They form the dorsal margin of naris anteriorly, the anterodorsal rim of the orbit posterolaterally, and contact the frontal and postorbital posteriorly (Figure 2A, B). The contact area between the prefrontal and frontal is convex forward, which is similar to *Nanhsiungchelys wuchingensis* (Tong & Li, 2019). In lateral view, the prefrontal is anterior to the postorbital and above the maxilla, and consists of the anterodorsal rims of the orbit (Figure 2C–F). This is similar to *Nanhsiungchelys wuchingensis*, *Jiangxichelys neimongolensis* and *Zangerlia ukhaachelys* (Brinkman & Peng, 1996; Joyce & Norell, 2005; Tong & Li, 2019). Behind the naris, the prefrontal is convex dorsally (Figure 2C–F), rather than concave downward showing as in *Nanhsiungchelys wuchingensis* (Tong & Li, 2019).

### Frontal

The frontal is a large pentagonal bone that is located in the center of the skull roof (Figure 2A, B), which is similar to *Nanhsiungchelys wuchingensis* and *Zangerlia ukhaachelys* (Joyce & Norell, 2005; Tong & Li, 2019). Its anterior margin has a “Λ” shape for articulating with the prefrontal. The lateral and posterior margins contact the postorbital and parietal respectively. It is excluded from the rim of the orbit, as in *Nanhsiungchelys wuchingensis* and *Zangerlia ukhaachelys* (Joyce & Norell, 2005; Tong & Li, 2019). There is no suture between the paired frontals, suggesting they are fused at the midline. This is an autapomorphy rather than ontogenetic variation, because the sutures occurred on the mature individual of *Nanhsiungchelys wuchingensis* (IVPP V3106) whose carapace length is 111 cm (Tong & Li, 2019).

### Postorbital

The postorbital is subtriangular in outline and elongated anteroposteriorly, and it consists of part of the lateral skull roof. Most parts of the postorbital are behind the orbit, but the anterodorsal process extends to the dorsal edge of the orbit (Figure 2C–F). Thus, the postorbital consists of the posterior-upper and posterior rims of the orbits, which is similar to *Nanhsiungchelys wuchingensis*, *Jiangxichelys ganzhouensis* and *Zangerlia ukhaachelys* (Joyce & Norell, 2005; Tong et al., 2016; Tong & Li, 2019). The postorbital contacts the prefrontal and frontal anteriorly, the jugal and quadratojugal ventrally, and the parietal medially (Figure 2A–F). In dorsal view, the shape of the posterior margin of the postorbital is uncertain due to its poor preservation and because it is partly obscured by the carapace. Besides, it is uncertain if the postorbital consists of the rim of temporal emargination. Notably, the postorbital in both *Nanhsiungchelys yangi* and *Nanhsiungchelys wuchingensis* are relatively large in size (Tong & Li, 2019), whereas just a small element forms the ‘postorbital bar’ in *Jiangxichelys ganzhouensis* and *Zangerlia ukhaachelys* (Joyce & Norell, 2005; Tong et al., 2016).

### Parietal

The trapezoidal parietal contributes to the posterior part of the skull roof (Figure 2A, B), which is similar to *Nanhsiungchelys wuchingensis* (Tong & Li, 2019). However, the paired parietals are bigger than the fused frontal in dorsal view, contrasting with *Nanhsiungchelys wuchingensis* (Tong & Li, 2019). The parietal contacts the frontal anteriorly and contacts the postorbital laterally, and these boundaries are not straight. Posteriorly, the parietal constitutes the upper temporal emarginations, but the absence of the posterior ends (especially the right part) of the parietal limits the identification of the rim of upper temporal emarginations.

### Mandible

The mandible is preserved in situ and tightly closed with the skull (Figure 2C–F). The location of the mandible is posterior and interior to the maxillae (Figure 3). Therefore, the beak is hidden, but the lower parts of the mandible can be observed. The symphysis is fused, which is similar to *Nanhsiungchelys wuchingensis* (Tong & Li, 2019). In ventral view, the most anterior part of the mandible appears slender, but the middle and posterior parts are robust (Figure 3), which differs from *Nanhsiungchelys wuchingensis* in which nearly all parts of the mandible are equal in width (Tong & Li, 2019).

### Carapace

Only the anterior parts of the carapace are preserved (Figure 2A, B). The preserved parts indicate there is a deep nuchal emargination and a pair of anterolateral processes, which are similar to those of *Anomalochelys angulata*, *Nanhsiungchelys wuchingensis* and *Nanhsiungchelys* sp. (SNHM 1558) (Hirayama et al., 2001; Hirayama et al., 2009; Tong & Li, 2019). In contrast, the carapaces of other species of nanhsiungchelyids (including *Basilemys*, *Hanbogdemys*, *Kharakhutulia*, *Jiangxichelys*, *Zangerlia*) usually have a shallow nuchal emargination and/or lack the distinctive anterolateral processes (Mlynarski, 1972; Sukhanov, 2000; Sukhanov et al., 2008; Tong & Mo, 2010; Danilov et al., 2013; Mallon & Brinkman, 2018). In dorsal view, each anterolateral process of *Nanhsiungchelys yangi* is very wide (nearly 90°), similar to *Nanhsiungchelys wuchingensis*; however, the anterolateral processes of *Anomalochelys angulata* and *Nanhsiungchelys* sp. (SNHM 1558) are crescent-shaped and horn-shaped respectively, both of which are sharper than *Nanhsiungchelys yangi* (Hirayama et al., 2001; Hirayama et al., 2009). Among the above species of *Nanhsiungchelys* and *Anomalochelys*, there are always a distinct protrusion at the tip of each anterolateral process, and this protrusion becomes more prominent on *Anomalochelys angulata* and *Nanhsiungchelys* sp. (SNHM 1558) (Hirayama et al., 2001; Hirayama et al., 2009). Besides, in *Nanhsiungchelys wuchingensis* and *Anomalochelys angulata*, the most anterior end of the process shows varying degrees of bifurcation (Hirayama et al., 2001; Tong & Li, 2019), but this bifurcation does not occur in *Nanhsiungchelys yangi* and *Nanhsiungchelys* sp. (SNHM 1558) (Hirayama et al., 2009). Due to the lack of sutures preserved on the surface of the carapace, it is difficult to determine whether these processes are composed of nuchal or peripheral plates. However, considering the similarity in shape of the anterolateral processes in *Nanhsiungchelys yangi* and *Nanhsiungchelys wuchingensis*, the anterolateral processes of *Nanhsiungchelys yangi* may be formed by the first peripheral plates (the same condition in *Nanhsiungchelys wuchingensis*).

### Plastron

A large plate under the mandible is identified as the anterior part of plastron (Figure 3). Despite there being little damage, the anterior edge of the epiplastron extends forward beyond the deepest part of nuchal emargination (Figure 3), similar to *Basilemys*, *Hanbogdemys*, *Jiangxichelys*, *Nanhsiungchelys*, and *Zangerlia* (Sukhanov, 2000; Danilov et al., 2013; Brinkman et al., 2015; Tong et al., 2016; Mallon & Brinkman, 2018; Tong & Li, 2019). The anterior part of the epiplastron is very thin, but the plates become thickened posteriorly and laterally (Figure 1). Although preserved poorly, the angle between the left and right edges is about 55°, which is wider than *Hanbogdemys orientalis* (Sukhanov, 2000). The epiplastra are paired and connected at the midline. Because only the anterior part of the entoplastron is preserved, it is hard to recognize its shape. The anterior edges of the entoplastron are very convex, and lead into the posterior part of the epiplastra. The angle between the two anterior edges (>110°) is larger than in *Nanhsiungchelys wuchingensis* (~100°) (Tong & Li, 2019). The identifiable scutes are only gular and humeral. In many nanhsiungchelyids, like *Basilemys praeclara*, *Basilemys morrinensis*, *Jiangxichelys ganzhouensis*, *Jiangxichelys neimongolensis*, *Hanbogdemys orientalis*, *Zangerlia dzamynchondi*, *Kharakhutulia kalandadzei* (Brinkman & Nicholls, 1993; Brinkman & Peng, 1996; Sukhanov, 2000; Sukhanov et al., 2008; Danilov et al., 2013; Tong et al., 2016; Mallon & Brinkman, 2018), there are usually intergular or extragular scutes beside the gular scutes, but this does not occur in *Nanhsiungchelys wuchingensis* (Tong & Li, 2019) and *Nanhsiungchelys yangi*. Moreover, the location and shape of the sulcus of *Nanhsiungchelys yangi* are similar to *Nanhsiungchelys wuchingensis* (Tong & Li, 2019). In *Nanhsiungchelys yangi*, the sulcus between the gular and humeral scutes can be identified and they are slightly curved and extend onto the entoplastron, which is similar to *Jiangxichelys neimongolensis* and *Nanhsiungchelys wuchingensis* (Brinkman & Peng, 1996; Brinkman et al., 2015; Tong & Li, 2019). However, in the other nanhsiungchelyids (e.g. *Kharakhutulia kalandadzei*, *Zangerlia dzamynchondi*, *Hanbogdemys orientalis*, *Yuchelys nanyangensis* and *Jiangxichelys ganzhouensis*), this sulcus is tangential to (or separated from) the entoplastron (Sukhanov, 2000; Sukhanov et al., 2008; Tong et al., 2012; Danilov et al., 2013; Tong et al., 2016).

## Discussion

### Taxonomy

Through comparison with a complete specimen (IVPP V3106) of *Nanhsiungchelys wuchingensis*, the large skull (length = 13 cm) of CUGW VH108 is inferred to correspond to a ~55.5 cm carapace length. This large body size, coupled with the special network of sculptures on the surface of the skull and shell, clearly demonstrates that CUGW VH108 belongs to Nanhsiungchelyidae (Li & Tong, 2017). Moreover, CUGW VH108 has laterally thickened epiplastron (Figure 1) and the anterior edge of the epiplastron extends forward of the deepest part of nuchal emargination (Figure 3), additional features that are diagnostic of Nanhsiungchelyidae (Li & Tong, 2017).

Within Nanhsiungchelyidae, CUGW VH108 differs from *Basilemys*, *Hanbogdemys*, *Kharakhutulia*, *Yuchelys*, and *Zangerlia* because all of these taxa have weak nuchal emargination and/or lack distinct anterolateral processes (Mlynarski, 1972; Sukhanov, 2000; Sukhanov et al., 2008; Tong et al., 2012; Danilov et al., 2013; Mallon & Brinkman, 2018). Moreover, CUGW VH108 differs from *Jiangxichelys ganzhouensis* and *Jiangxichelys neimongolensis* in which the cheek emargination and temporal emargination are deep (Brinkman & Peng, 1996; Tong et al., 2016). Although the carapace of both *Anomalochelys* and CUGW VH108 have deep nuchal emargination and a pair of anterolateral processes, the former’s anterolateral processes are crescent-shaped and have a bifurcated anterior end (Hirayama et al., 2001), which are significant differences from the processes of CUGW VH108.

CUGW VH108 can be assigned to the genus *Nanhsiungchelys* because of the deep nuchal emargination, pair of anterolateral processes, and the weakly developed cheek emargination and temporal emargination (Li & Tong, 2017). However, CUGW VH108 differs from *Nanhsiungchelys wuchingensis* in which the snout is trumpet shaped (Tong & Li, 2019). Moreover, *Nanhsiungchelys wuchingensis* and CUGW VH108 have some different skeletal features on the skull, the two most important of which are that CUGW VH108’s premaxilla is very small and higher than wider (Figure 2C–F) and that a small portion of the maxilla extends behind and below the orbit (Figure 2C–F). CUGW VH108 also differs from *Nanhsiungchelys* sp. (SNHM 1558) in which the anterolateral processes are horn-shaped (Hirayama et al., 2009). Thus, CUGW VH108 differs from all other known species of Nanhsiungchelyidae, and herein we erect a new species *Nanhsiungchelys yangi*. *Nanhsiungchelys yangi* is a medium-sized species of *Nanhsiungchelys*, with an estimated carapace length of more than 0.5 meters. The surface of the skull, carapace, and plastron are covered in a special network of sculptures consisting of pits and ridges. The triangular-shaped snout is large and long. Both the cheek emargination and temporal emargination are weakly developed. The carapace has a deep nuchal emargination and a pair of wide anterolateral processes.

### Phylogenetic position and paleobiogeography

The phylogenetic analysis retrieved seven most parsimonious trees with a length of 77 steps, with a consistency index (CI) of 0.675 and retention index (RI) of 0.679. The strict consensus tree (Figure 8) recovers *Nanhsiungchelys yangi* and *Nanhsiungchelys wuchingensis* as sister taxa, and one unambiguous synapomorphy was identified: the absence of the extragulars. These two species and *Anomalochelys angulata* form a monophyletic group, which is consistent with the results of Tong & Li (2019). Synapomorphies of this group include that the neurals are wide, the anterior side of the first vertebral is constricted and primarily in contact with the cervical only, and the ratio of length to width of the carapace is larger than 1.6. In particular, our new character (character 50, the ratio of length to width of the carapace) also effectively proves their relationship, which further suggests the ratio of carapace needs more attention in turtles’ phylogeny. However, the standard bootstrap and Bremer supports values are low among these groups, and their relationships therefore need further consideration. Interestingly, our new results identify *Yuchelys nanyangensis* and *Zangerlia testudinimorpha* as sister taxa, and this relationship was supported by one unambiguous synapomorphy (their fifth vertebral almost fully covers the suprapygal).

Our results support a close relationship between the genera *Nanhsiungchelys* and *Anomalochelys*, even though they lived in different times and regions (Figure 8). However, based on the similarity of extinct plants and animals, Sun & Yang (2010) inferred that the Japan Sea did not exist during the Jurassic and Cretaceous, with the Japan archipelago still closely linked to the eastern continental margin of East Asia. This view is also supported by geological and geophysical evidence (Kaneoka et al., 1990; Liu et al., 2017). In addition to *Anomalochelys angulata* from Hokkaido (Hirayama et al., 2001), many fragments of Nanhsiungchelyidae (as *Basilemys* sp.) have also been found on Honshu and Kyushu islands, Japan (Hirayama, 1998, 2002; Danilov & Syromyatnikova, 2008). In China, the easternmost specimen of nanhsiungchelyids (a fragment of the shell) was recovered from the Upper Cretaceous of Laiyang, Shandong (Li & Tong, 2017), which is near the west coast of the Pacific Ocean and close to Japan geographically. This geographical proximity likely allowed nanhsiungchelyids to migrate between China and Japan during the Late Cretaceous.

*Nanhsiungchelys* is the only group of turtles that has been found from the Upper Cretaceous of Nanxiong Basin, and it shows a high diversity, including *Nanhsiungchelys yangi*, *Nanhsiungchelys wuchingensis*, and *Nanhsiungchelys* sp (SNHM 1558) (Hirayama et al., 2009; Tong & Li, 2019). Such high diversity in a restricted space is comparable to the condition on some other islands, for example, 14 species of *Chelonoidis* lived on nine islands of the Galapagos archipelago (Zhou & Zhou, 2020). Perhaps the reason for this phenomenon is that *Nanhsiungchelys* was adapt to the extreme environment of Nanxiong Basin, such as the hot temperatures (26.66~33.95 °C) during the Late Cretaceous (Yang et al., 1993), with the constituent species occupying different ecological niches.

### Function of the anterolateral processes of the carapace

*Nanhsiungchelys* and *Anomalochelys* have a pair of distinct anterolateral processes on the carapace, and we hypothesized that these processes would affect drag when the animal was swimming through water. The CFD simulations allow us to evaluate the drag produced by each of the turtle models. In simulations with flow velocities ranging from 0.6 m/s to 2.6 m/s, both the drag forces and the drag coefficients of the *Nanhsiungchelys yangi* model were always lower than the generalized turtle model (Table 3; Figure 9). Considering the only difference between these two 3-D models is whether there is a pair of anterolateral processes on carapace, this result strongly suggests that these processes played an important role in reducing resistance (~25 % reduction in drag). The reduction of drag could enhance locomotory performance by conserving the energy expended during swimming (Fish, 2000; Gutarra et al., 2019; Song et al., 2021). This reinforces the importance of the anterolateral processes to the movement of *Nanhsiungchelys* in water.

**Table 3.** Drag forces and drag coefficients for three-dimensional digital models of *Nanhsiungchelys yangi* and a generalized turtle at different flow velocities.

| Flow velocity<br>(m/s) | *Nanhsiungchelys yangi* |  | Generalized turtle |  | Drag reduction due to<br>anterolateral processes |
| --- | --- | --- | --- | --- | --- |
|  | Drag force (N) | Drag coefficient | Drag force (N) | Drag coefficient |  |
| 0.6 | 7.132 | 0.330 | 9.473 | 0.439 | 24.71 % |
| 1.0 | 19.176 | 0.320 | 25.826 | 0.430 | 25.75 % |
| 1.4 | 37.064 | 0.315 | 50.204 | 0.427 | 26.17 % |
| 1.8 | 60.734 | 0.312 | 82.586 | 0.425 | 26.46 % |
| 2.2 | 90.204 | 0.311 | 122.988 | 0.424 | 26.66 % |
| 2.6 | 125.444 | 0.309 | 171.410 | 0.423 | 26.82 % |

In the simulations with an inlet flow velocity of 1.0 m/s, the 2-D plots of flow velocity magnitude show that there was a low-velocity zone near the proximal forelimbs in the generalized turtle model (Figure 10A), but this zone was not evident in the *Nanhsiungchelys yangi* model (Figure 10B). The same pattern was observed when the inlet flow velocity was increased to 2.2 m/s (Figure 10C, D). The reason for this phenomenon may be that the anterolateral processes of *Nanhsiungchelys yangi* did not extend horizontally forward, but instead bend downwards. Therefore, these processes made the anterior part of the shell more streamlined (Figure 11A, B), analogous to the streamlined fairing on the anterior of airplanes and rockets. In the model without processes, the anterior edge of the shell is rather blunt, resulting in greater overall drag (Figure 11C, D).

Hirayama et al. (2001) assumed that the horns (processes) on *Anomalochelys*’ carapace were used to protect the skull. However, due to the lack of obvious neural crest and transverse processes on the cervical vertebra, Yeh (1966) suggested that the neck of *Nanhsiungchelys wuchingensis* was flexible, and inferred that the skull could be withdrawn into the shell to avoid danger. *Jiangxichelys neimongolensis* was likely also able to withdraw the head into the shell, as indicated by a complete specimen (93NMBY-2) from Inner Mongolia, China (Brinkman et al., 2015). Thus, if nanhsiungchelyids were able to protect the skull and neck by withdrawing them into the shell, the anterolateral processes may have had a different function. The anterolateral processes are unlikely to have been used for mate competition. In extant tortoises (e.g. *Testudo horsfieldi*), males compete for mates by hitting each other with the anterior edges of their plastron (Shi, 1998), and this is why male tortoises usually have more robust anterior edges of plastron. However, it is unknown whether nanhsiungchelyids similarly fought for the right to mate, and the prominent processes on *Nanhsiungchelys* and *Anomalochelys* are on their carapace rather than the plastron. Thus, a role in enhancing locomotion through drag reduction is the most likely explanation for the presence of the anterolateral processes on the carapace.

The swimming ability of *Nanhsiungchelys* is not incompatible with some skeletal characteristics of terrestrial turtles (Yeh, 1966). This is parallel with some extant tortoises (e.g. *Aldabrachelys gigantea*), which has a domed carapace and elephantine limbs, like burying the body in the mud of shallow water to avoid the hot weather (Zhou & Zhou, 2020), and they could even swim (or float) in the ocean (Gerlach et al., 2006; Hansen et al., 2016). Perhaps the drag-reducing function of the anterolateral processes of *Nanhsiungchelys* helps them survive in a harsh environment and even migrate over a long distance.

## Conclusions

A turtle skeleton (CUGW VH108) with a well-preserved skull and lower jaw, together with the anterior parts of the shell, was found in Nanxiong Basin, China. This is assigned to the genus *Nanhsiungchelys* based on the huge estimated body size (~55.5 cm), a special network of sculptures on the surface of the skull and shell, weak cheek emargination and temporal emargination, deep nuchal emargination, and a pair of anterolateral processes on the carapace. Based on the character combination of a triangular-shaped snout and wide anterolateral processes, we erect a new species *Nanhsiungchelys yangi*. A phylogenetic analysis of nanhsiungchelyids places *Nanhsiungchelys yangi* and *Nanhsiungchelys wuchingensis* as sister taxa. *Nanhsiungchelys* shows a high diversity (three species) in Nanxiong Basin, which may be because these turtles could adapt to extremely hot environments during the Late Cretaceous. Finally, based on the results of CFD, we infer that the anterolateral processes on the carapace in *Nanhsiungchelys yangi* could enhance locomotory performance by reducing drag when the animal was swimming through water.

## Supporting information

Appendix 1. Taxon-character matrix (nex.)

Appendix 2. Three-dimensional digital models of Nanhsiungchelys yangi and generalized turtle (stl.)

Appendix 3. Reconstruction steps of three-dimensional digital models

## Acknowledgements

We thank Xing Xu (IVPP) for his useful suggestions, thank Kaifeng Wu (YSNHM) for preparing turtle skeleton, and thank Mingbo Wang (UPC), Zichuan Qin (UB), Wen Deng (CUGW) and Haoran Sun (LJU) for assistance with CFD. This work was supported by the National Natural Science Foundation of China (42288201).

## Appendix 1. Taxon-character matrix (nex.)

## Appendix 2. Three-dimensional digital models of *Nanhsiungchelys yangi* and generalized turtle (stl.)

## Appendix 3. Reconstruction steps of three-dimensional digital models

## Notes

### Competing Interest Statement

The authors have declared no competing interest.

