## Appendix 3. Reconstruction steps of three-dimensional digital models for "A new species of *Nanhsiungchelys* (Testudines: Cryptodira: Nanhsiungchelyidae) from the Upper Cretaceous of Nanxiong Basin, China, and the role of anterolateral processes on the carapace in drag reduction"

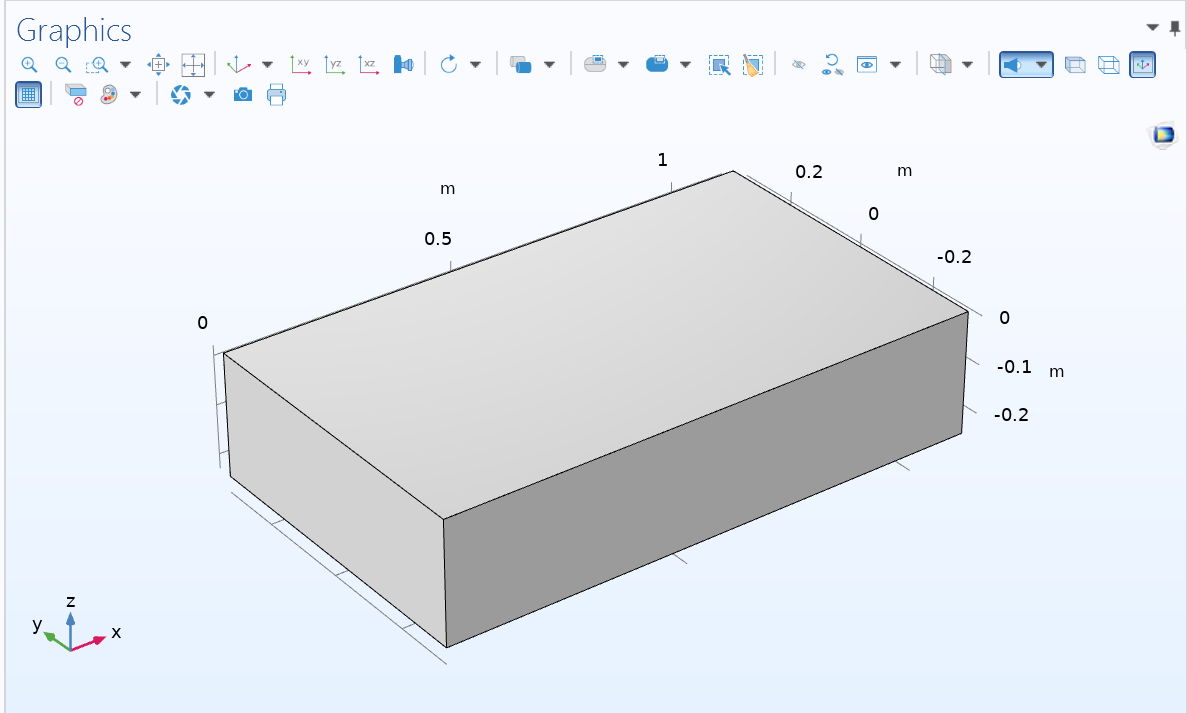

（1） Build a block (blk1).

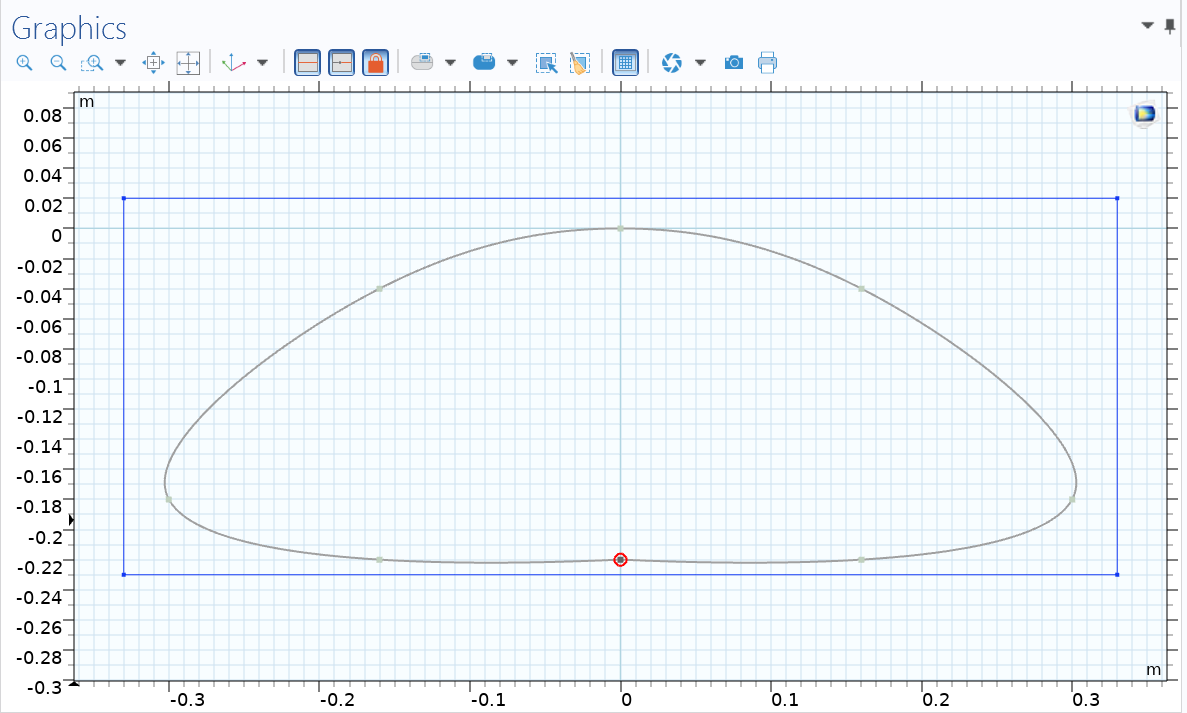

(2) Establishing a work plane (wp1) along Y-Z direction, and building an interpolation curve.

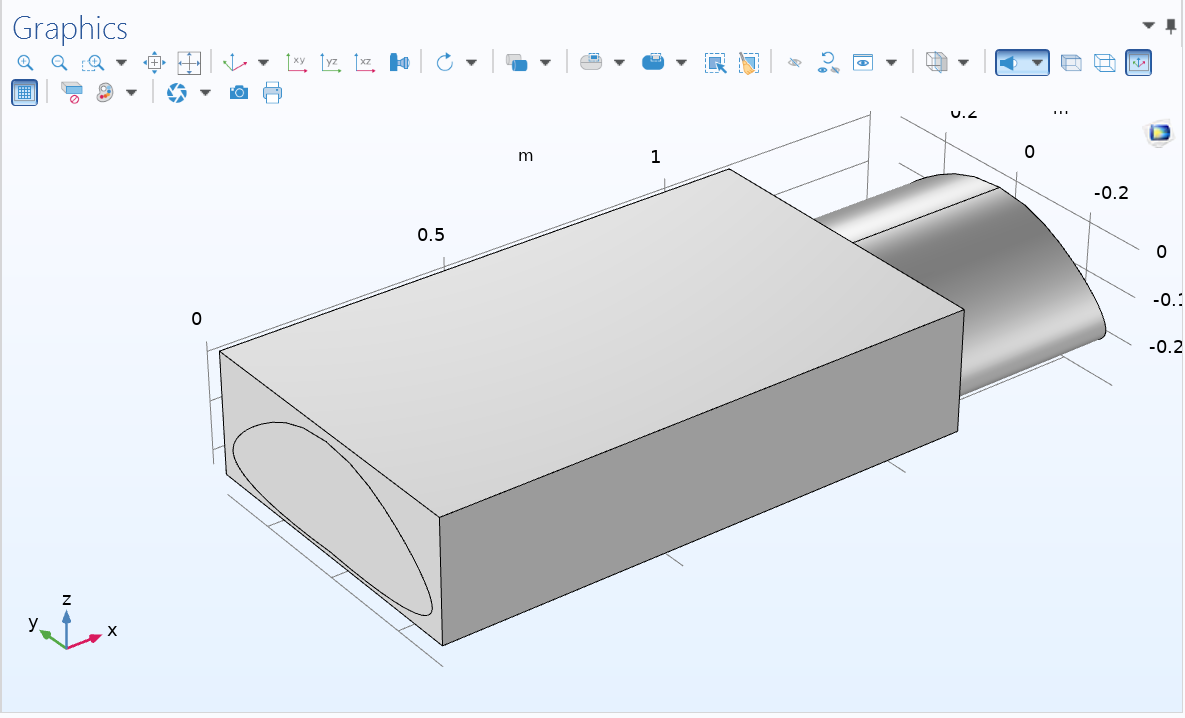

(3) Extend the interpolation curve to the X-axis direction to form the extended plane (ext1).

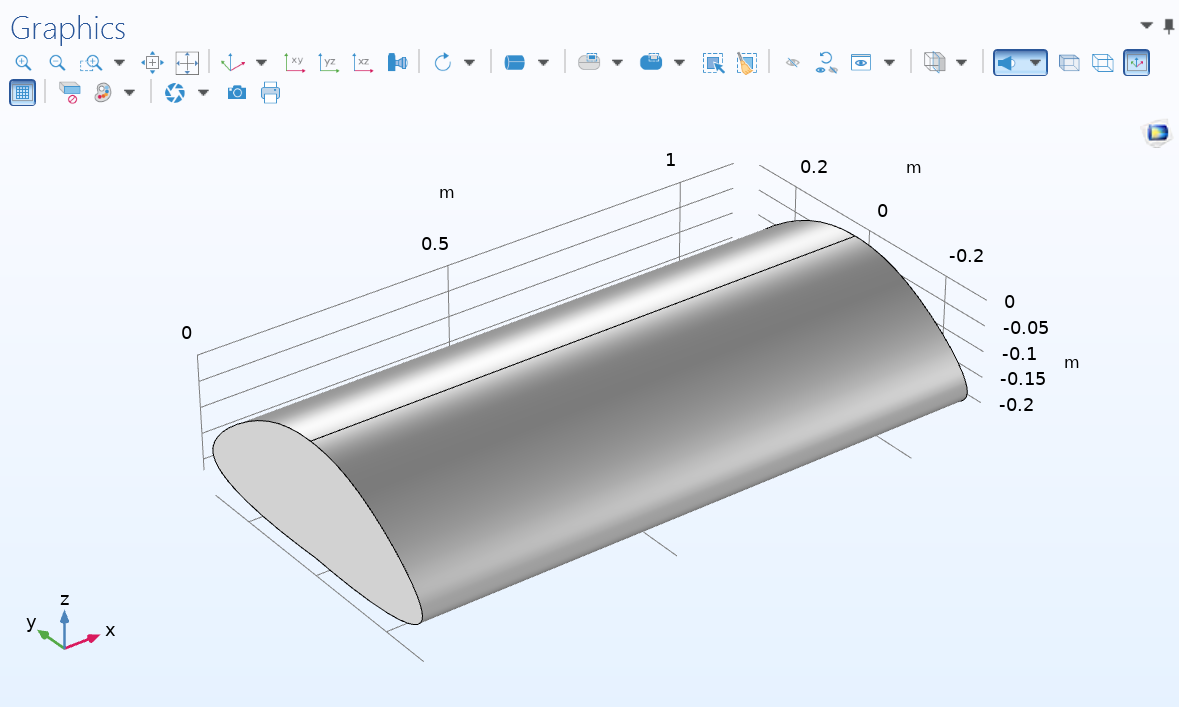

(4) Partitioning the block (blk1) with ext1. Then, delete ext1 and redundant domains.

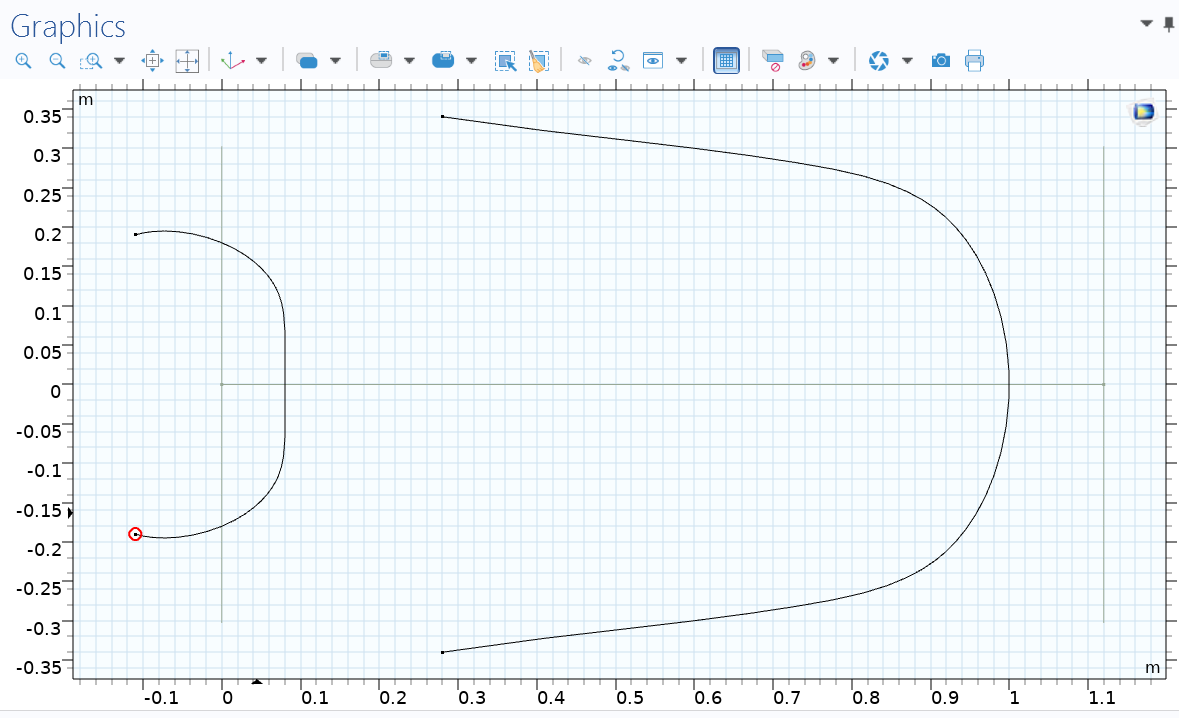

(5) Establishing another work plane (wp2) along X-Y direction, and building an interpolation curve.

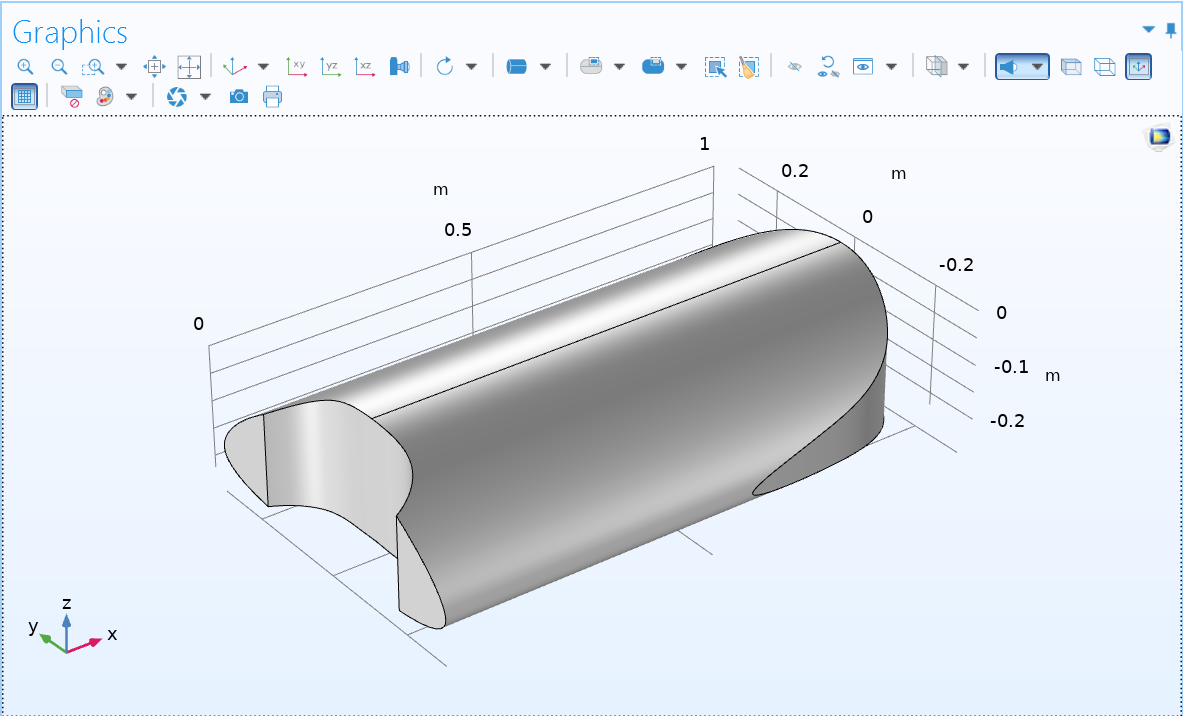

(6) Building an extended plane (ext2). Partitioning the domain with ext2. Then, delete ext2 and redundant domains. The detailed methods are similar to steps 3 and 4.

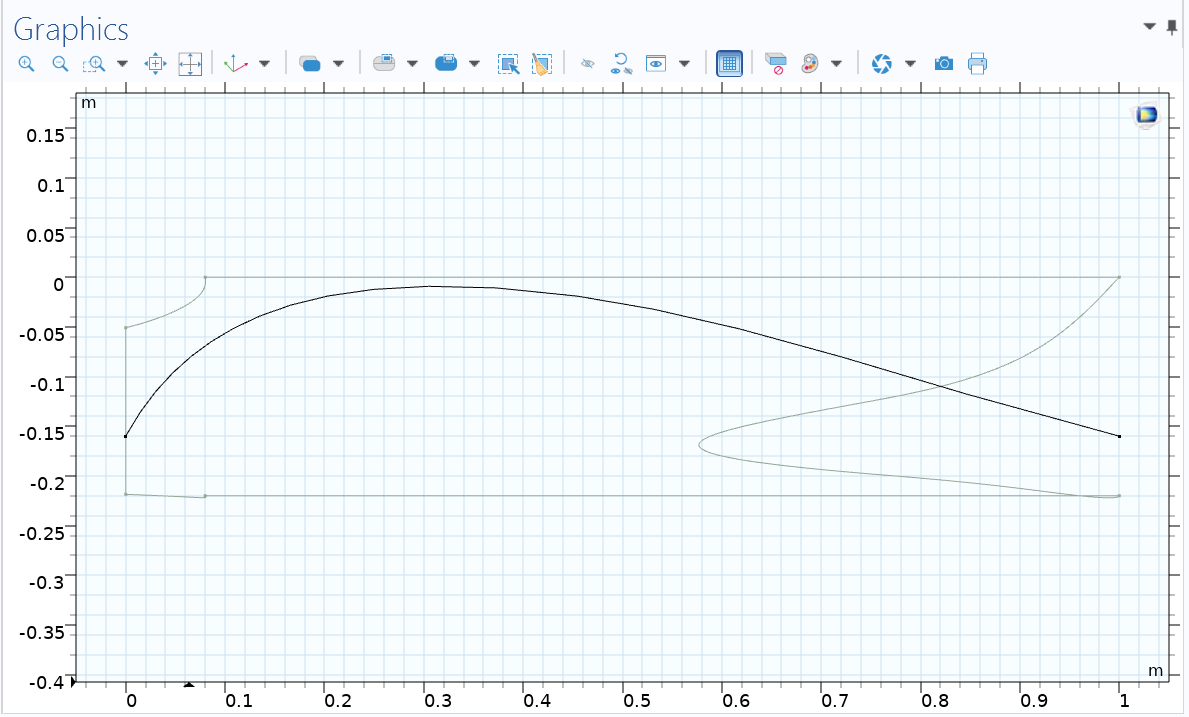

(7) Establishing a new work plane (wp3) along X-Z direction, and building an interpolation curve.

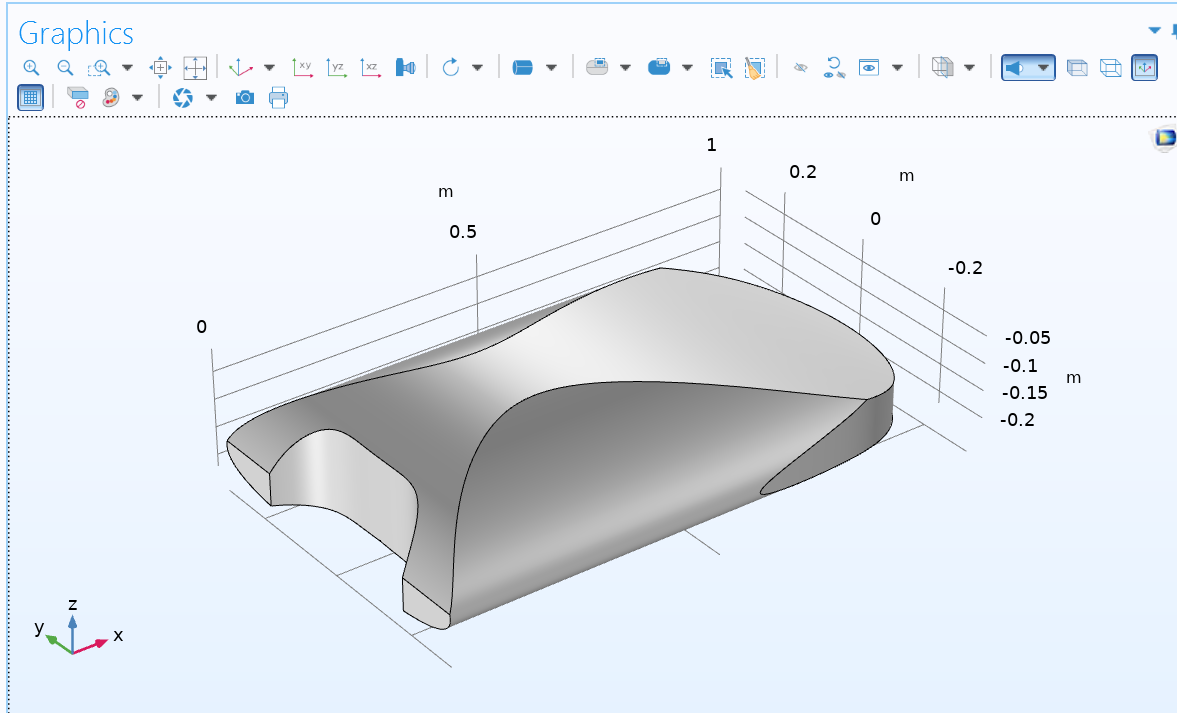

(8) Building an extended plane (ext3). Partitioning the domain with ext3. Then, delete ext3 and redundant domains. The detailed methods are similar to steps 3 and 4.

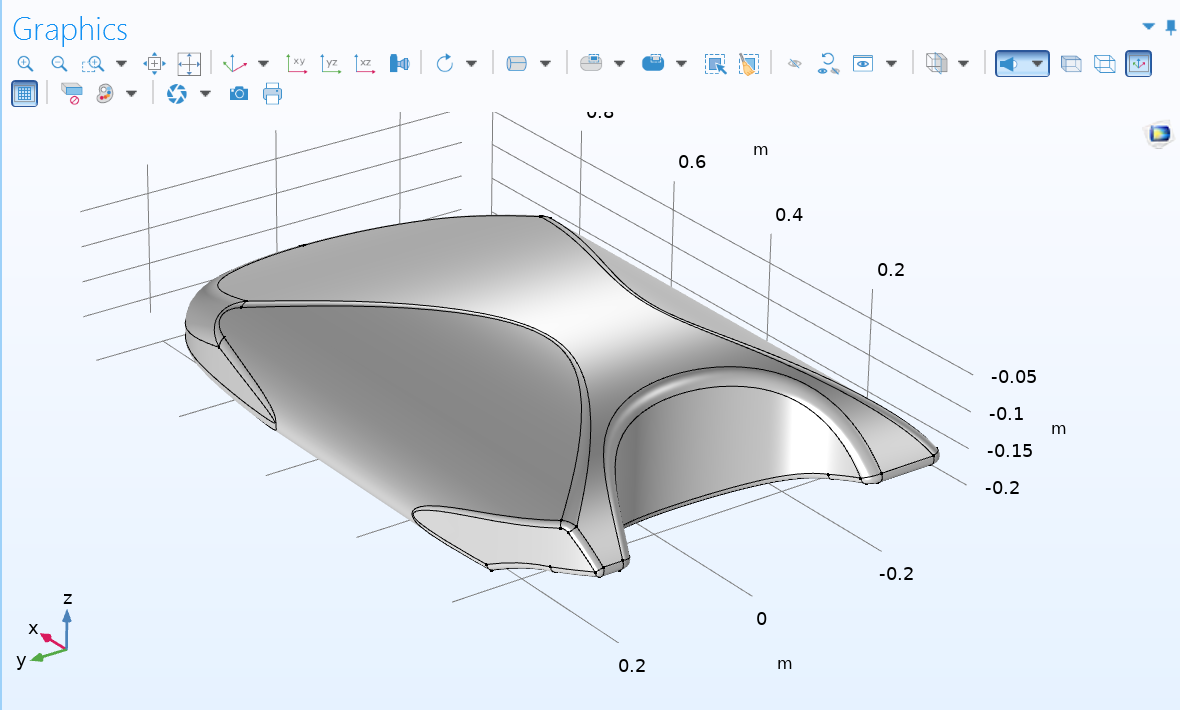

(9) It is necessary to improve some details. For example, the anterolateral processes and nuchal emargination could also be modified using the previous methods. Moreover, some edges could be smooth using the Fillet function. Of course, we can modify only one half (as shown in the figure), and then use symmetry for the other half.

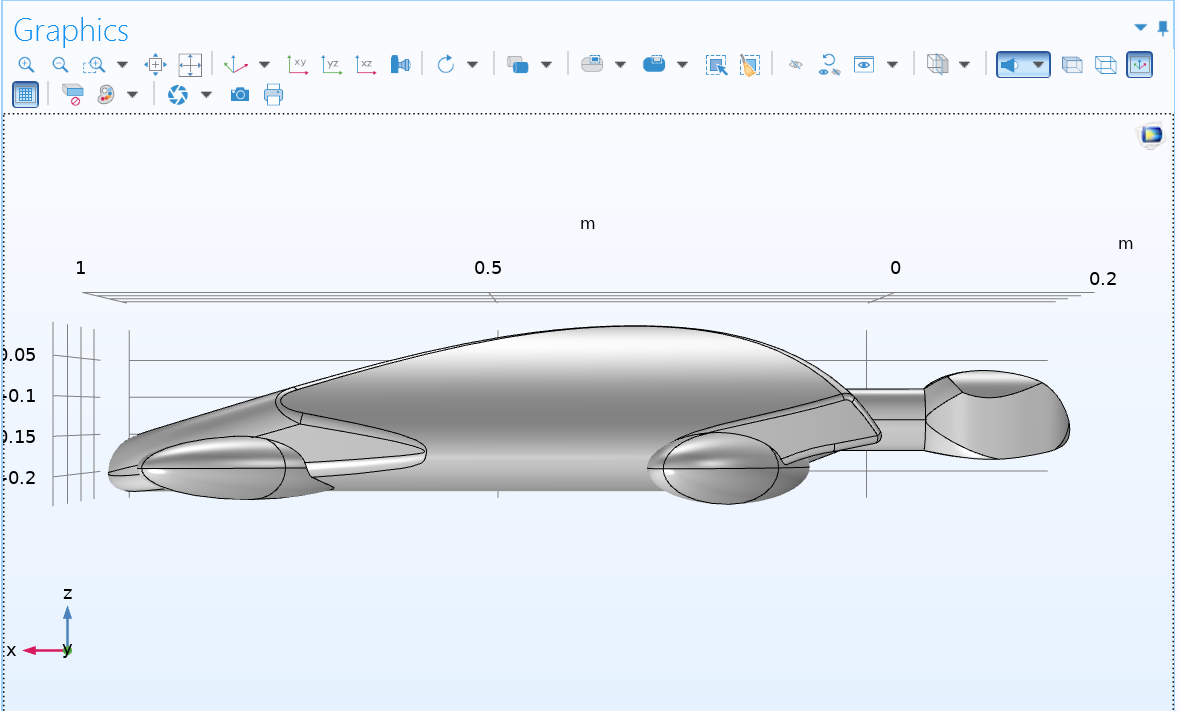

(10) Building the head, neck, and limbs, which are composed of some geometry (e.g. ellipsoids, cylinders). The redundant domains are partitioned and deleted.

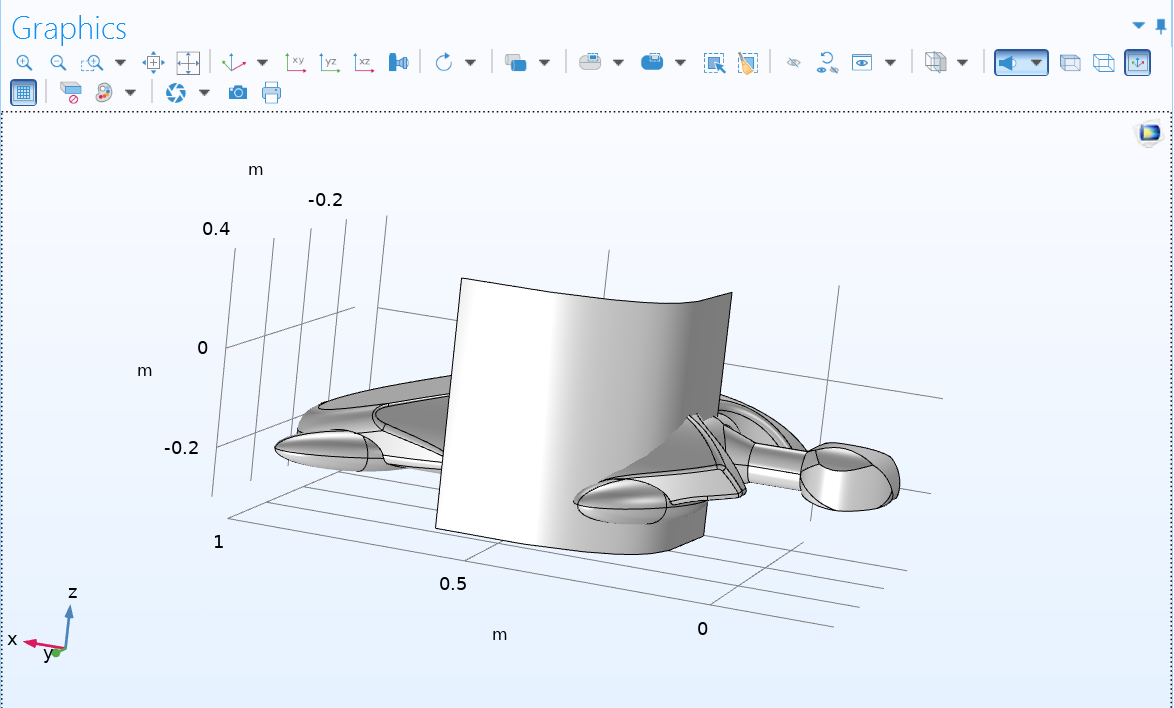

(11) If cut off the anterolateral processes, a generalized turtle model could be produced. This could also be done by building an extended plane.
